# Genome Assembly of the Green Revolution Wheat Cultivar Pavon 76 Establishes a Reference for CIMMYT-Derived Wheat

**DOI:** 10.64898/2026.06.04.730269

**Authors:** Ethan Eurmsirilerd, Mariano Resendiz, Giuseppe Lana, Qing Li, Fatima Nawmi, Rendell Chang, Wei Zhang, Timothy X. Wu, Zuoming Sun, Lili Wang, Stefano Lonardi, Joanna Melonek, John M. Chater, Zhenyu Jia

## Abstract

Common wheat (*Triticum aestivum* L.) is a cornerstone of global food security. However, the standard reference genome, IWGSC RefSeq v2.1, is derived from wheat cultivar Chinese Spring, a historically important lineage that is not representative of modern cultivated wheat. In contrast, Pavon 76 is a CIMMYT Green Revolution wheat cultivar grown worldwide and is present in the pedigrees of numerous elite wheats, as well as in the genetic background of over 600 cytogenetic stocks used in studies ranging from recombination to male sterility. As no Pavon 76 reference has been reported, interpretation of these Pavon 76-derived stocks has relied on the genetically distant IWGSC RefSeq v2.1. To address this gap, we generated PacBio HiFi reads and Hi-C read pairs to assemble Pavon 76’s genome. The final assembly comprises the expected 21 chromosomes, spans 15 Gb, and has an N50 of 710 Mb. Annotation identified 167,201 genes, of which 89.5% were functionally annotated. Comparative sequence and gene synteny analysis between Pavon 76 and IWGSC RefSeq v2.1 revealed significant structural divergence, establishing that IWGSC RefSeq v2.1 is not fully representative of modern wheats. This robust Pavon 76 assembly establishes the first chromosome-scale *de novo* assembly of a CIMMYT-derived wheat and provides a foundation for future genetic and genomic studies.

## 1. Introduction

Common wheat (*Triticum aestivum* L.) is a cornerstone of global food security and remains one of the world’s most important staple crops (Shewry & Hey, 2015). Given its global agricultural importance, wheat has been the focus of extensive cytogenetic and genomic research, resulting in numerous assembled genomes representing elite cultivars. The current standard wheat reference genome, IWGSC RefSeq v2.1, is derived from cv. Chinese Spring, an old wheat cultivar from China whose precise origin and pedigree remain unknown (International Wheat Genome Sequencing Consortium [IWGSC], 2018; Liu *et al*., 2018; Zhu *et al*., 2021). Chinese Spring became the wheat genetic standard largely through the development of an extensive set of cytogenetic stocks created by Dr. Ernest Robert Sears of the United States Department of Agriculture and the University of Missouri (Sears, 1988). Chinese Spring’s selection for these genomic studies was largely historical, originating from a fortuitous discovery of two haploids after a failed attempt at chromosome doubling of wheat-rye hybrids (Sears, 1988). While the IWGSC RefSeq v2.1 represented a landmark achievement in wheat genomics and advanced the study of wheat genetics and cytogenetics, Chinese Spring is not representative of modern cultivated wheat. Consequently, reliance on the IWGSC RefSeq v2.1 limits the characterization of structural and functional variation present in globally important elite germplasm. Despite the widespread global use of germplasm developed by the International Maize and Wheat Improvement Center (CIMMYT, El Batán, Mexico), no chromosome-scale, high-quality *de novo* reference genome has been reported for any CIMMYT-derived wheat cultivar, leaving a major gap between available genome assemblies and the cultivars that dominate global agriculture (Liu *et al*., 2026).

In contrast to Chinese Spring, Pavon 76 (hereinafter Pavon) is a representative Green Revolution wheat cultivar developed by CIMMYT and widely grown for its broad environmental adaptability and agronomic performance. Pavon is a hard white spring wheat, with the pedigree VICAM 71//CIANO 67’S’/SIETE CERROS 66/3/KALYANSONA/BLUEBIRD, which includes the classics of the Green Revolution (URGI, n.d.). Its developer, Dr. Sanjaya Rajaram, Director of the Global Wheat Program at CIMMYT, considered it a “global wheat” because of its demonstrated performance across diverse environments (personal communication). Pavon’s strong performance led to its widespread use in breeding programs, and it is present in the pedigrees of numerous wheat cultivars, including hard red winter wheats of the U.S. Great Plains (Nebraska Wheat, 2018). In addition to its importance in breeding, Pavon has served for decades as a foundational stock for over 600 cytogenetic and genetic stocks. The genetic stocks include chromatin introgressions from related species such as rye (*Secale cereale*), *Agropyron elongatum, Haynaldia villosa,* and others, as well as numerous aneuploid stocks and chromosome structural variants (Adam J. Lukaszewski, personal communication). These materials have been widely used in studies of chromosome pairing control, recombination, male sterility, and the control of double fertilization in wheat (Fan *et al*., 2021; Fan *et al*., 2024; Lukaszewski & Brzezinski, 2003; Lukaszewski, 2024). However, interpretation of these materials has historically relied on the genetically distant Chinese Spring reference genome, limiting the accurate characterization of cultivar-specific structural variation. A *de novo* Pavon reference genome assembled independently of existing wheat references would therefore provide an essential genomic framework for interpreting these genetic stocks.

Historically, wheat genome assembly has been challenging due to its size and complexity. Bread wheat is an allopolyploid, composed of three sub-genomes (AABBDD) originating from ancestral diploid progenitors: *Triticum urartu* (sub-genome A), a presumably extinct relative of *Aegilops speltoides* (sub-genome B), and *A. tauschii* (sub-genome D) (Levy & Feldman, 2022). Extensive homoeology among corresponding chromosomes and repetitive content of approximately 85% have historically complicated high-quality genome assembly. Recent advances in Pacific Biosciences High-Fidelity (HiFi) long-read sequencing and Hi-C scaffolding have substantially improved the assembly of complex polyploid genomes and enabled chromosome-scale wheat assemblies with high contiguity (Mascher *et al*., 2021; Sato *et al*., 2021).

In this study, we integrated 30-fold HiFi coverage with approximately 33-fold Arima Genomics Hi-C data to produce a complete *de novo* chromosome-level genome assembly of Pavon. The resulting assembly resolves the expected 21 single-chromosome pseudomolecules, with high contiguity, addressing the lack of a CIMMYT-derived wheat reference genome and filling a major gap in wheat genomic resources.

## 2. Materials and Methods

### 2.1 DNA Extraction and Genome Sequencing

High-molecular-weight DNA for PacBio HiFi sequencing was extracted from fresh tissue of wheat cv. Pavon seedlings that were germinated and then grown in the absence of light, using a modified Cetrimonium bromide (CTAB) protocol that excels at the extraction of high molecular weight DNA (Resendiz *et al*., 2025).

HiFi libraries for the PacBio Revio SMRT Cell long-read sequencing were prepared at the University of California, Riverside Genomics Core Facility. The DNA was size-selected using the PacBio Short Read Eliminator (SRE) XL kit, then slightly fragmented to around 15-25 kb using the g-TUBE (Covaris), and cleaned by AmPure XP beads (Beckman Coulter). The quality and quantity of the resulting HMW DNA were measured using a 4150 TapeStation System (Agilent Technologies) with a concentration of 117 ng/µL and a size distribution of 7-25 kbp. The DNA libraries were made using the PacBio HiFi Prep Kit according to the manufacturer’s protocol, and their quality was assessed using Femto Pulse Systems (Agilent Technologies). Sequencing was performed on a PacBio Revio sequencer at the University of California, Davis. A total of six SMRT Cells were used, with a >1800.02 movie time and a 120-minute pre-extension time. The raw data were processed using the CCS version 8.2.0. HiFi reads were constructed when subreads with nine or more passes were obtained in each cell. The CCS analysis report summary statistics for each SMRT Cell are available in Supplemental Table 1.

For the Arima Genomics Hi-C data, fresh tissue from the same seedlings as the PacBio HiFi sequencing was frozen in liquid nitrogen and shipped on dry ice to Arima Genomics (Carlsbad, CA). Hi-C data were generated using the Arima High Coverage HiC kit, according to the manufacturer’s protocols. Sequencing was performed on a NovaSeq X (Illumina) with 150 bp paired-end reads on a 25B flow cell lane, yielding 1,812,690,954 read pairs.

### 2.2 Sequencing Data Quality Assessment

To assess the quality of the PacBio HiFi reads, LongQC v1.2.1 was used with default parameters on each of the six PacBio SMRT Cell FASTQ files (Fukasawa *et al*., 2020). The six files were then combined into a single FASTQ file and used with Jellyfish v2.3.0 to create a histogram of k-mer frequencies with k = 81 (Marçais & Kingsford, 2011). GenomeScope2 v2.0.1 then used the Jellyfish histogram to estimate the k-mer coverage (Ranallo-Benavidez *et al*., 2020). To assess for any potential contamination from other species during sequencing, the six FASTQ files were analyzed with Kraken 2 v2.0.8 (Wood *et al*., 2019).

Arima Genomics prepared Hi-C libraries for Pavon and, in parallel, created a positive control (GM12878 cell line control library), which was sequenced. The positive control sequencing data were then analyzed using the Arima-SV pipeline, which revealed a high-quality library with low PCR duplicates and a high proportion of long-range cis contacts and low trans contacts. Following this step, the Pavon library was sequenced at a high depth. Quality control of the paired-end reads was carried out using FASTQC v0.11.8 (https://www.bioinformatics.babraham.ac.uk/projects/fastqc/).

### 2.3 De novo Chromosome-Scale Nuclear Genome Assembly

A *de novo* assembly was generated from 481.11 Gb of HiFi reads (roughly a 30-fold coverage) using hifiasm v0.25.0 (Cheng *et al*., 2021). As the Pavon line was highly inbred, purging during genome assembly or analysis was neither necessary nor used. We used the primary assembly from hifiasm for all subsequent steps. Following this, Arima Hi-C read pairs were mapped to the hifiasm assembly using the Arima Mapping Pipeline v03 (<u>https://</u> <u>github.com/ArimaGenomics/mapping_pipeline</u>) with default parameters. The scaffolded assembly was generated with the Yet Another Hi-C Scaffolding tool v1.2.2 (YaHS) using default parameters, integrating the assembly from hifiasm and the Arima Mapping Pipeline output (Zhou *et al*., 2023). Gaps in the scaffolded assembly were then filled using the raw HiFi reads with TGS-Gapcloser v1.2.1 (Xu *et al*., 2020). The scaffolded assembly was corrected using PretextViewAI v1.0.1 and the Rapid Curation Pipeline to correct orientation errors and misjoins (https://gitlab.com/wtsi-grit/rapid-curation). Chromosome naming and orientation were determined by mapping Pavon pseudomolecules to the IWGSC RefSeq v2.1 with minimap2 v2.30 (Li, 2018).

### 2.4 De Novo Organelle Genome Assembly and Annotation

*De novo* assembly of the organelle genomes was performed using Oatk v1.0 using all HiFi reads generated (Zhou *et al*., 2025). Annotation of the organelles was performed using GeSeq v2.03 (Tillich *et al*., 2017). For the mitochondrial genome, *T. aestivum* cvs. Chinese Spring (MH051716.1) and Yumai (EU534409.1), *T. timopheevii* (NC_022714.1), and *Amblyopyrum muticum* (OZ209253.1) reference genomes were used in GeSeq. For the plastid genome, *T. aestivum* (KC912694) and the specific cultivar Chinese Spring TA3008 (KJ614396), *T. timopheevii* (AB976560.1), and *A. muticum* (PQ632086) were used. Organellar Genome DRAW (OGDRAW) was used to produce annotation figures (Grenier *et al*., 2019). Unplaced contigs aligning to organelles were removed from the final nuclear assembly.

### 2.5 Nuclear Genome Quality Assessment

The final assembly was evaluated with QUAST v5.3.0 using default parameters, with IWGSC RefSeq v2.1 as the reference for assembly statistics (Gurevich *et al*., 2013). In addition, BUSCO v5.8.2 in “genome-mode” evaluated genome completeness using the poales_odb12 dataset and default parameters (Tegenfeldt *et al*., 2025). Hi-C contact maps of the pseudomolecules were generated using HiCExplorer v3.6 with a bin size of 1,000,000 bp following the pipeline at https://hicexplorer.readthedocs.io/en/latest/content/example_usage.html (Wolff *et al*., 2020). To measure HiFi read coverage, reads were aligned to the finished pseudomolecules using minimap2 v2.30. The alignment was used as input for NucFreq v0.1, which computed nucleotide frequencies (Vollger *et al*., 2019). Two versions of NucFreq plots were created: one without a sequence read depth limitation (Supplemental Figure 1), and one with a sequence read depth limitation of 100x (Supplemental Figure 2).

### 2.6 Nuclear Genome Annotation

First, the final assembly was annotated using CLARI-TE v1.0 (https://forge.inrae.fr/umr-gdec/clari-te_smk), a pipeline specialized for transposon annotation in wheat (Daron *et al*., 2014). Subsequently, the EviAnn v2.0.4 pipeline was used to annotate the full nuclear genome by integrating numerous evidence sources (Zimin *et al*., 2025). Homology evidence was provided by Lifton v1.0.8, which transfers annotations from the IWGSC RefSeq v2.1 to the Pavon assembly (*Chao et al*., 2025). For *ab initio* gene prediction, ANNEVO v2.2.2 annotated the CLARI-TE hard-masked Pavon assembly using the Embryophyta model (Zhang *et al*., 2026). Protein evidence included sequences from the Ensembl plant databases of *T. durum*, *T. urartu*, *A. tauschii*, *Hordeum vulgare*, *Oryza sativa*, *Brachypodium distachyon*, *Zea mays*, and *Arabidopsis thaliana*, and Uniprot/SwissProt (Clade Poales, taxid: 38820). Transcript evidence consisted of publicly available Iso-Seq reads from 14 wheat tissue types (Liu *et al*., 2025), which were mapped with Minimap2 v2.30 and assembled with StringTie2 v3.0.2 (Kovaka *et al*., 2019). Summary statistics from the annotation were obtained using AGAT v1.4.0 (Dainat *et al*., 2026), and a circle plot of the annotation features was produced in R v4.5.0 using circlize v0.4.16 (Gu *et al*., 2014). Following structural annotation, functional annotation was performed using eggNOG-mapper v2.1.9 (Cantalapiedra *et al.,* 2021). Annotation quality was assessed by running BUSCO v5.8.2 on the translated protein sequences from EviAnn’s final annotation output, using the poales_odb12 dataset with default parameters.

### 2.7 Comparisons of Pavon 76 to IWGSC RefSeq v2.1

To compare the final Pavon assembly with the existing IWGSC RefSeq v2.1 reference, QUAST v5.3.0 was run on the Pavon (Supplemental File 1) and IWGSC RefSeq v2.1 (Supplemental File 2) pseudomolecules, and the resulting statistics were compared. Pavon was aligned to the IWGSC RefSeq v2.1 assembly using minimap2 v2.30. The resulting whole-genome alignments were used as input for Synteny and Rearrangement Identifier (SyRI) v1.7.1 to identify and visualize structural rearrangements between Pavon and IWGSC RefSeq v2.1 (Goel *et al*., 2019). To compare gene annotations between Pavon and IWGSC RefSeq v2.1, synteny analysis was run using GENESPACE v1.3.1 (Lovell *et al*., 2022).

## 3 Results

### 3.1 Chromosome-Scale Genome Assembly

A total of 40.1 million PacBio HiFi reads were generated, providing 30-fold genome coverage (Supplemental Table 1). The average N50 for the HiFi reads across all SMRT Cells was 12.01 kb. LongQC confirmed high read quality across all SMRT Cells, with a median read quality of Q39.5. GenomeScope2 k-mer depth matched the coverage, providing a 29.7-fold coverage depth (Figure 1A). GenomeScope2’s estimated monoploid genome size of 2.37 Gb corresponded to an estimated hexaploid genome size of 14.22 Gb, which is consistent with other wheat genome assemblies and nearly matched the size of the 21 pseudomolecules of this assembly. When checked for contamination, Kraken 2 found that across all SMRT Cells, over 95% of reads were assigned to *Triticum*, consistent with expectations. No contamination was detected.

**Figure 1.**
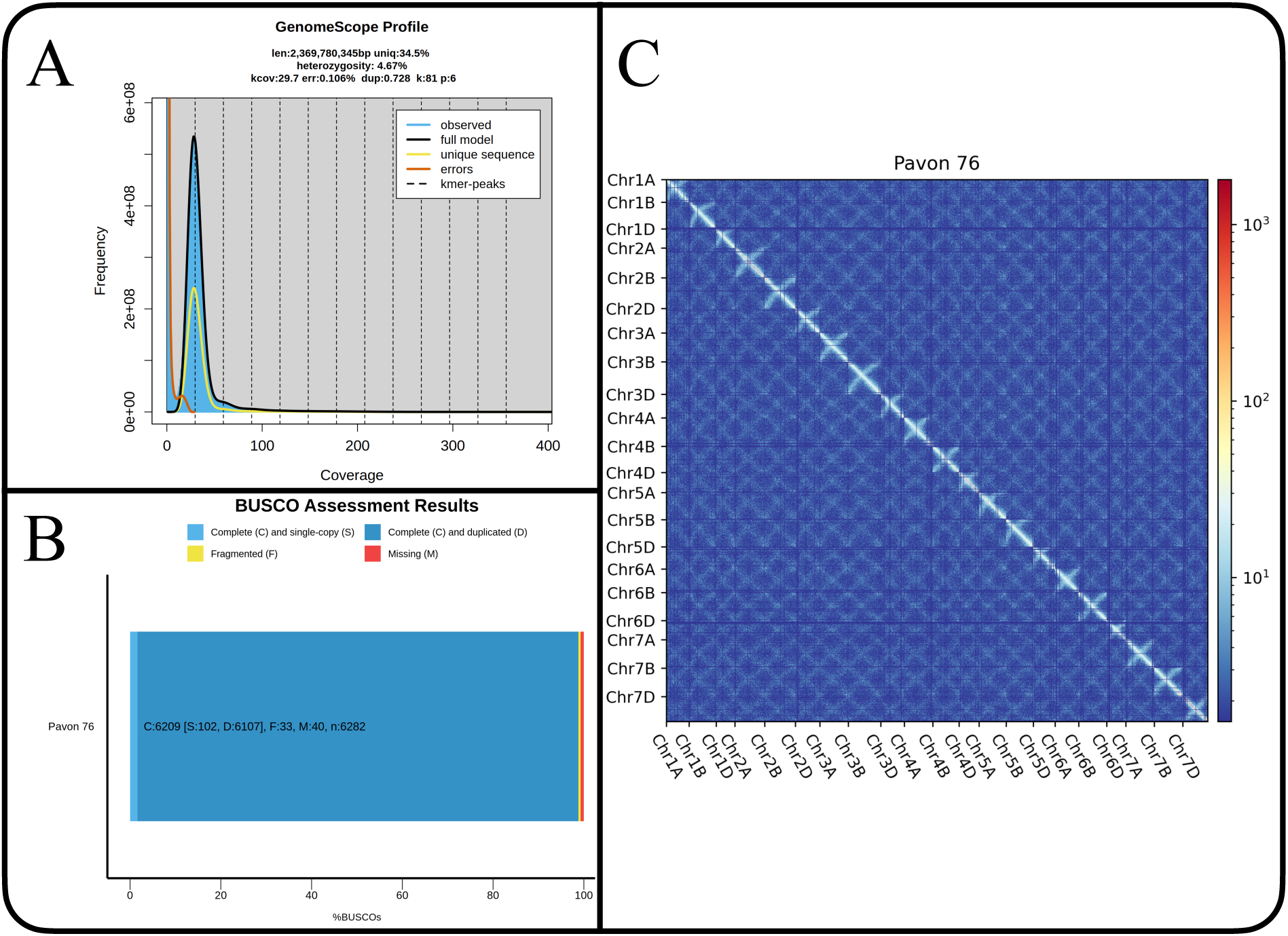
Quality Assessment of Pavon 76 Pseudomolecule Assembly. A: GenomeScope2 profile for 81-mers based on HiFi reads. B: BUSCO Assessment Result. C: Genome-wide Hi-C contact heat map with a bin size of 1 Mb.

The PacBio HiFi reads were assembled using hifiasm, and the k-mer frequency plot showed a single peak around 30, consistent with a homozygous sample at both the targeted sequencing coverage depth and the GenomeScope2 k-mer depth. Arima Hi-C reads were mapped to this assembly using the Arima Mapping Pipeline and used to create a scaffolded assembly with YaHS. Following gap closure with TGS-gapcloser, manual curation using the Rapid Curation pipeline corrected several misassemblies. The final assembly comprises all 21 chromosomes, each in a single-chromosome pseudomolecule, and 8,522 unplaced contigs, totaling 15.06 Gb and an N50 of 709.76 Mb (Table 1). Compared to reported wheat genome assemblies, these metrics are comparable to or exceed those of other high-quality chromosome-scale assemblies (Liu *et al*., 2026). Approximately 95% of the genome has been placed into one of the 21 single-chromosome pseudomolecules, which together span 14.25 Gb, comparable to the 14.23 Gb represented by the IWGSC RefSeq v2.1 pseudomolecules. The total length of the unplaced contigs is 815.82 Mb. Pavon pseudomolecule statistics are available in Table 2.

**Table 1.** Pavon 76 Assembly Summary Statistics.

|  | **Pavon 76 Assembly** |
| --- | --- |
| Total Length (bp) | 15,064,574,134 |
| Number of Pseudomolecules and Contigs | 8,543 |
| N50 (bp) | 709,756,138 |
| N90 (bp) | 504,481,683 |
| L90 (no.) | 20 |
| Largest Contig (bp) | 851,610,634 |
| Number of Ns per 100 kbp | 0.72 |
| GC Content (%) | 46.36 |

**Table 2.** Pavon 76 Pseudomolecule Statistics.

|  | Length (bp) | Number of Contigs | Number of Genes |
| --- | --- | --- | --- |
| Chr1A | 602,377,969 | 23 | 5,848 |
| Chr1B | 705,049,253 | 85 | 5,903 |
| Chr1D | 498,741,800 | 15 | 6,000 |
| Chr2A | 780,276,094 | 36 | 7,684 |
| Chr2B | 811,264,797 | 69 | 7,989 |
| Chr2D | 643,756,051 | 24 | 7,684 |
| Chr3A | 753,024,579 | 17 | 7,077 |
| Chr3B | 851,610,634 | 91 | 7,579 |
| Chr3D | 623,742,505 | 28 | 7,323 |
| Chr4A | 744,112,795 | 55 | 6,470 |
| Chr4B | 691,635,960 | 83 | 4,746 |
| Chr4D | 527,888,988 | 23 | 5,350 |
| Chr5A | 709,756,138 | 21 | 7,229 |
| Chr5B | 716,684,190 | 98 | 6,981 |
| Chr5D | 578,219,328 | 24 | 7,668 |
| Chr6A | 621,552,855 | 22 | 5,626 |
| Chr6B | 736,946,254 | 98 | 6,069 |
| Chr6D | 504,481,683 | 14 | 5,938 |
| Chr7A | 743,581,887 | 32 | 7,556 |
| Chr7B | 750,282,457 | 94 | 5,966 |
| Chr7D | 653,770,019 | 24 | 7,738 |
| Unanchored Contigs | 815,817,898 | 8,522 | 26,777 |

### 3.2 Nuclear Genome Assembly Quality Assessment

Regarding assembly completeness, over 98.8% of BUSCO genes were found as complete (Figure 1B). This finding matches or exceeds other wheat genomes, such as cv. Fielder (97.1%) and Julius (98.3%), which have high levels of completeness (Sato *et al*., 2021). As expected for an allopolyploid consisting of three parental genomes, cvs. Pavon (Figure 1B), Fielder, and Julius all have high duplication rates, exceeding 90% (Sato *et al*., 2021).

Genome integrity of the pseudomolecules was assessed using genome-wide Hi-C contact maps (Figure 1C) and pseudomolecule-level Hi-C contact maps (Figure 2). These did not indicate any misassemblies. NucFreq analysis revealed no major coverage irregularities outside highly repetitive telomeric or centromeric regions, and showed low heterozygosity consistent with a highly inbred, self-pollinating line (Supplemental Figures 1 and 2).

**Figure 2.**
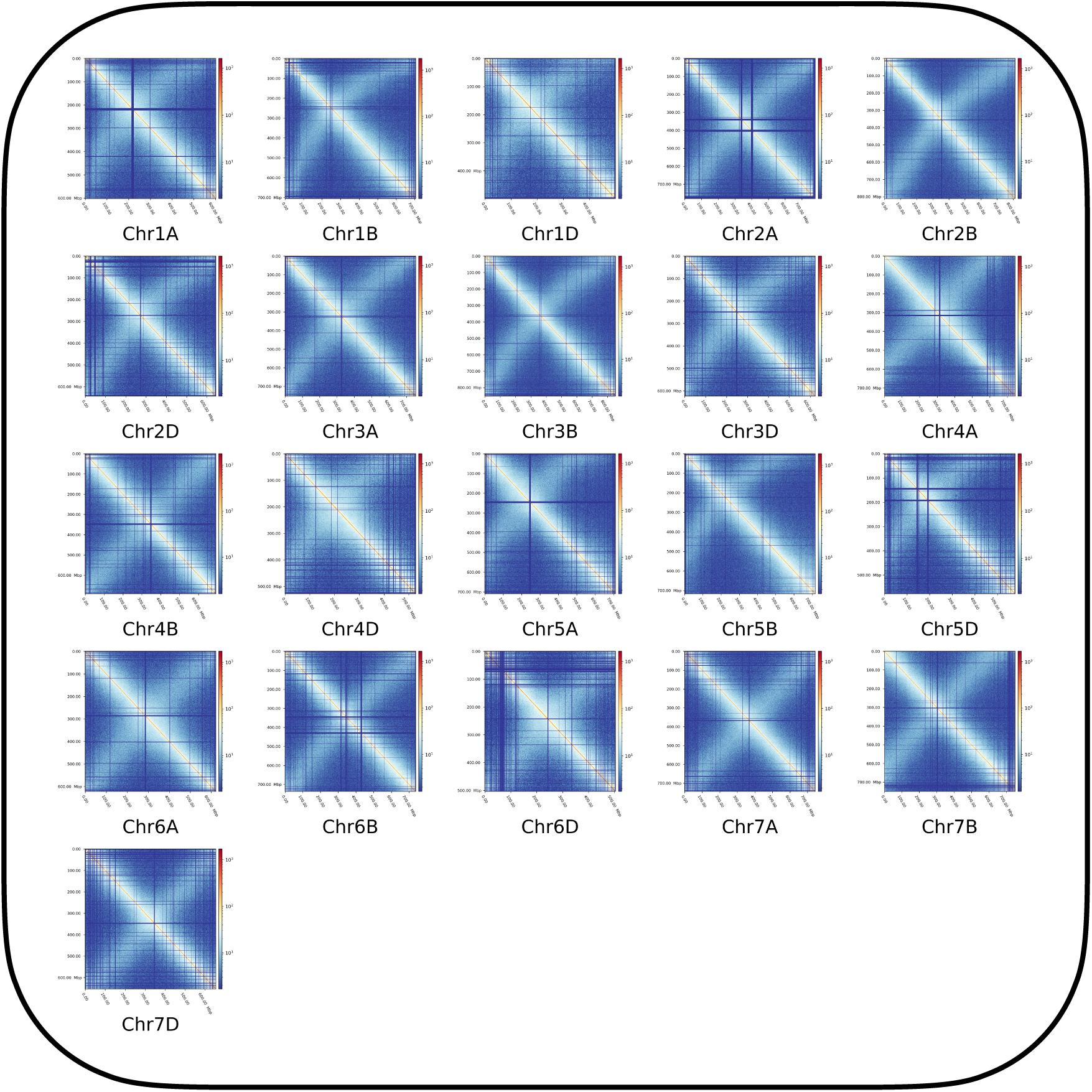
Pavon 76 pseudomolecule-level Hi-C contact maps with a bin size of 1 Mb.

### 3.3 Nuclear Genome Annotation

Overall, 83.5% of the genome was classified as transposable elements by Clari-TE. The final annotation identified 167,201 genes, of which 149,641 (89.5%) were functionally annotated. When the proteins in the annotations were tested with BUSCO, 99.6% of the BUSCO genes were identified as complete. The slightly higher completeness relative to the genome-based BUSCO analysis (98.8%) likely reflects differences between nucleotide- and protein-based assessments, with both supporting the assembly’s overall completeness. Summary statistics are presented in Table 3, and a circle plot showing gene density, transposable element density, and GC content across all pseudomolecules is shown in Figure 3. The circle plot showed transposable element enrichment in proximal chromosomal regions and higher gene density toward the distal chromosome ends, consistent with previous wheat genome analyses (Daron *et al*., 2014; Erayman *et al*., 2004).

**Table 3.** Pavon 76 Structural Annotation Statistics.

| **Statistic** | **Value** |
| --- | --- |
| Number of Genes | 167,201 |
| Number of Transcripts | 227,771 |
| Transcripts per Gene | 1.4 |
| Number of Monoexonic Genes | 68,645 |
| Number of Monoexonic Transcripts | 87,865 |
| Mean Transcript Length (bp) | 3,151 |
| Minimum Transcript Length (bp) | 14 |
| Maximum Transcript Length (bp) | 785,975 |
| Total Number of Exons | 984,511 |
| Mean Number of Exons per Transcript | 4.3 |
| Mean Exon Length (bp) | 322 |
| Mean CDS Length (bp) | 1,160 |
| Mean Protein Length (aa) | 348.9 |
| Number of Introns | 756,740 |

| Statistic | Value |
| --- | --- |
| Mean Intron Length (bp) | 528 |
| Mean 5'UTR Length (bp) | 275 |
| Mean 3' UTR Length (bp) | 476 |
| Total Exon Length (bp) | 317,574,977 |
| Total Intron Length (bp) | 400,216,869 |
| Total Gene Length (bp) | 484,647,995 |

**Figure 3.**
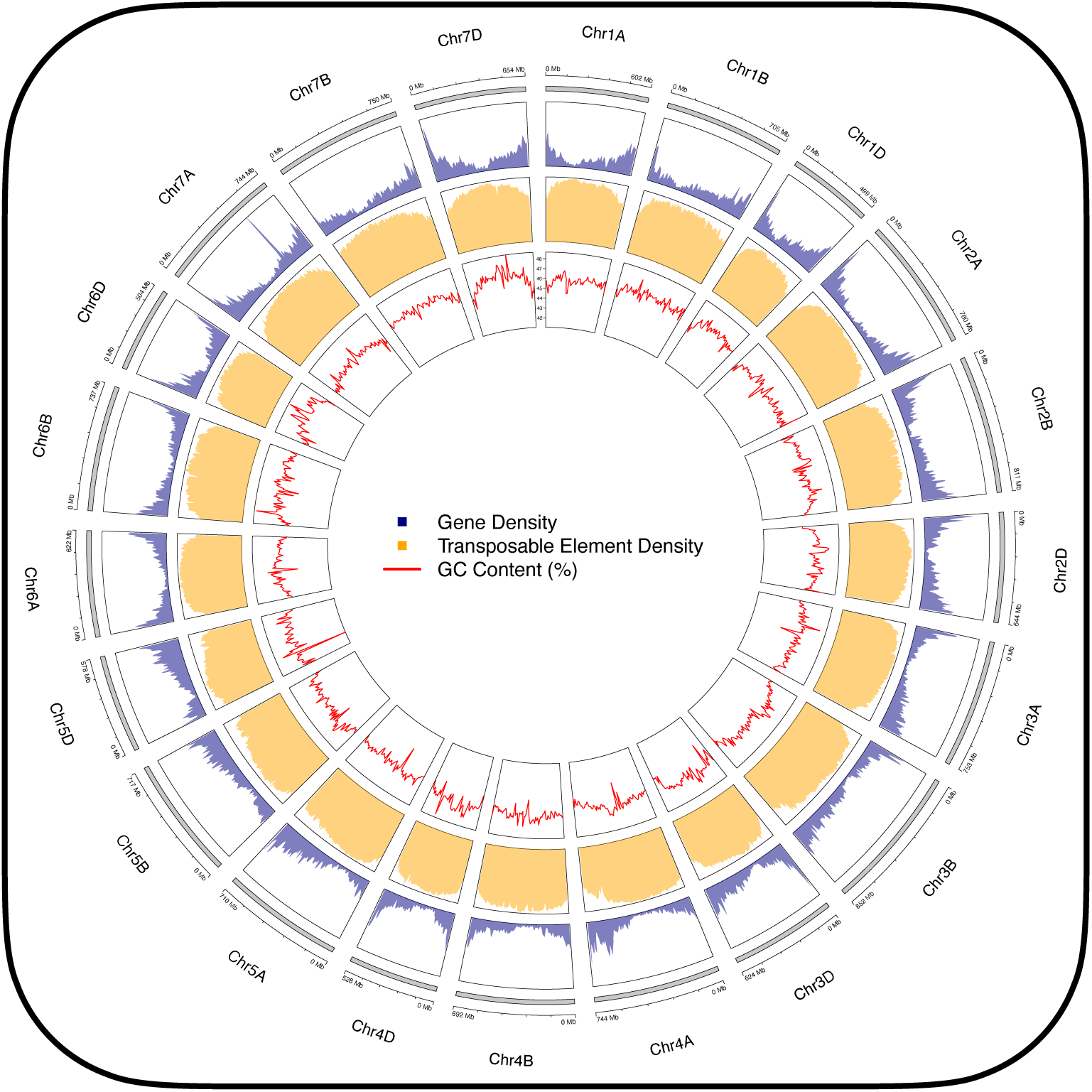
Circle plot of Pavon 76 assembly showing chromosome length, gene density, transposable element density, and GC content. The rings are, from outermost to innermost, chromosome length, gene density, transposable element density, and GC content (%).

### 3.4 De Novo Organelle Genome Assembly and Annotation

The complete mitochondrial and chloroplast genomes of Pavon 76 were obtained using the high-throughput long-read HiFi data generated for the nuclear genome assembly. The high accuracy and length of HiFi reads enabled the mitochondrial and chloroplast genomes to be assembled into complete circular sequences. The assembled Pavon 76 mitogenome was 452,522 bp in length (Figure 4), matching the sizes previously reported for *T. aestivum* (AABBDD) (Ogihara *et al*., 2005; Cui *et al*., 2009), but was slightly shorter than that of Hoh501 *T. turgidum* (AABB) and longer than that of *T. urartu* (AA) (Hu *et al*., 2023). Annotation and sequence analysis allowed us to identify a total set of 53 genes to be encoded in the mitochondrial genome (Supplemental Table 2), including 34 protein-coding genes, 3 rRNA genes, and 16 tRNA genes. Among these, one or two additional partial sequences were identified for 12 mitochondrial protein-coding genes, whereas four partial sequences were found for the *atp1* gene. Such partial sequences may contribute to the generation of novel open reading frames (ORFs) through mitochondrial genome recombination events, and reflect the dynamic nature of plant mitochondrial genomes.

**Figure 4.**
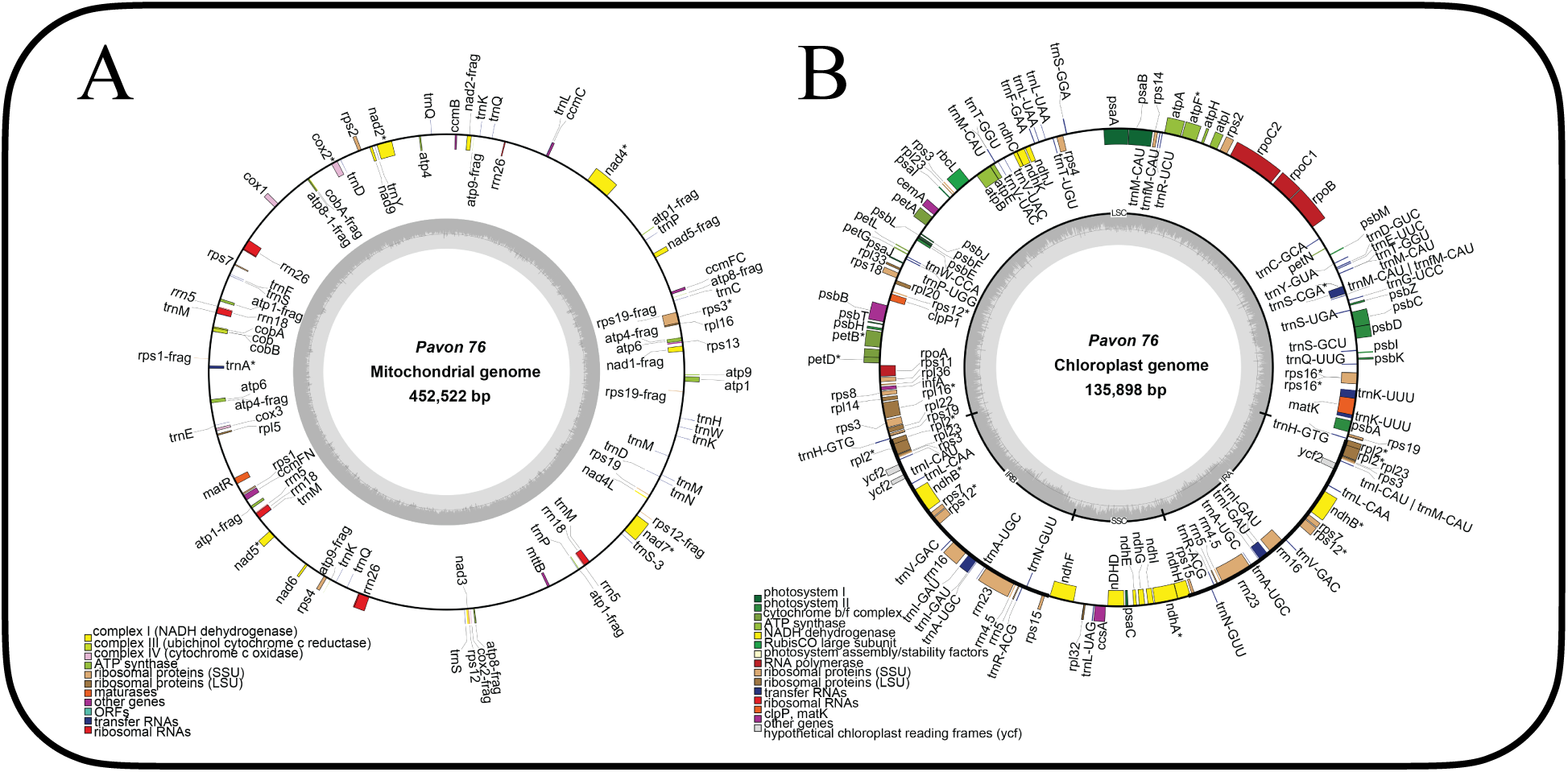
Pavon 76 Organellar Genome DRAW plot of the organelle assemblies. A: Mitochondria. B: Chloroplast. Abbreviation: frag, fragment.

Similar to most wheat chloroplast genomes, the chloroplast genome of Pavon 76 exhibited a typical quadripartite structure, consisting of a large single-copy (LSC) region, a small single-copy (SSC) region, and two inverted repeat regions (IRa and IRb) (Figure 4). The complete chloroplast genome of Pavon 76 was 135,898 bp, with the LSC, SSC, IRa, and IRb regions measuring 81,289 bp, 13,733 bp, 20,438 bp, and 20,438 bp, respectively (Figure 4). The chloroplast genome of Pavon 76 contained 110 genes in total, which fell into four main functional categories: photosynthesis (44), expression-related genes (59), genes of unknown function (3), and others (4) (Supplemental Table 3). A total of 27 genes were present in multiple copies. Among them, some genes, such as *ndhB*, *rps3*, and *rps7*, were represented by two identical copies, whereas *trnA-UGC* and *trnM-CAU* were present in four and six copies, respectively. Most chloroplast protein-coding genes were highly conserved, with sequence identities generally around 99% compared with the wheat and wheat wild relative chloroplast reference genomes used for annotation (Gornicki *et al*., 2014; Middleton *et al*., 2014; Zhang *et al*., 2025).

### 3.5 Comparisons to IWGSC RefSeq v2.1

Comparisons of the pseudomolecules of Pavon and the IWGSC RefSeq v2.1, performed using QUAST, revealed similarity across nearly all metrics, except for an over 2,000-fold difference in the N’s per 100 kb, with Pavon having 0.72 compared to 1524.94 in IWGSC RefSeq v2.1 (Supplemental Files 1 and 2). Despite comparable chromosome-level contiguity, this disparity reflects the improved resolution of long-read sequencing in Pavon, relative to earlier short-read-based assemblies such as the IWGSC RefSeq v2.1. SyRI analysis based on whole-genome alignments identified high levels of overall synteny, alongside substantial structural variation comprising 20,670 events spanning 6.05% of the genome (Table 4). These findings highlight the notable structural divergence between Pavon 76 and the IWGSC RefSeq v2.1, consistent with their origins (Figure 5).

**Table 4.** Rearrangements between Pavon 76 and IWGSC RefSeq v2.1

|  | Number of Events | Cumulative Span (bp) | % of the Genome |
| --- | --- | --- | --- |
| Inversion | 482 | 767,740,870 | 5.10% |
| Translocation | 3,550 | 65,049,016 | 0.43% |
| Duplication | 16,638 | 79,022,298 | 0.52% |
| Total | 20,670 | 911,812,184 | 6.05% |

**Figure 5.**
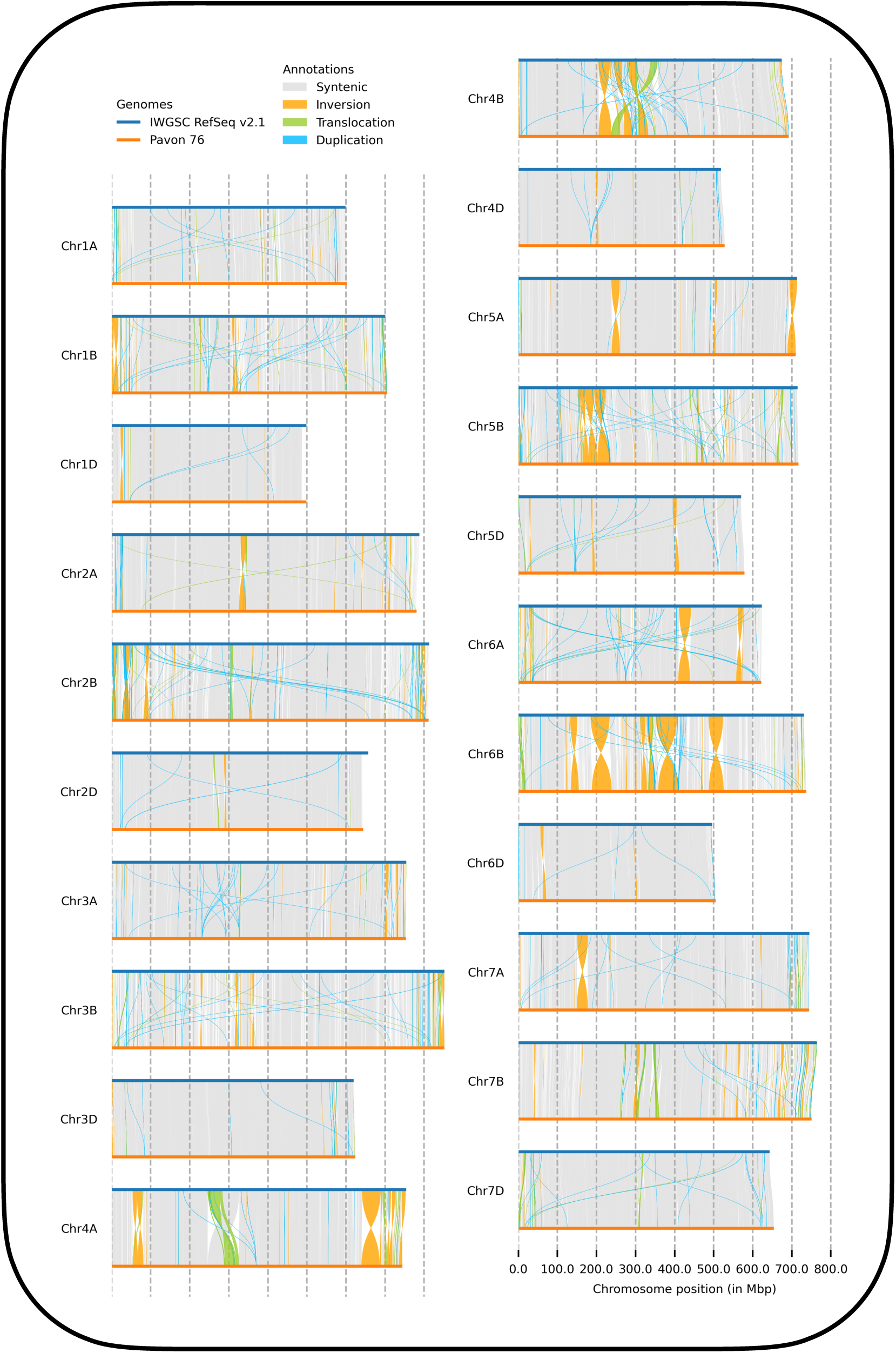
Synteny and Rearrangement Identifier (SyRI) results comparing the Pavon 76 and IWGSC RefSeq v2.1 (cv. Chinese Spring) assemblies.

Synteny analysis of the Pavon and IWGSC RefSeq v2.1 annotations was conducted using GeneSpace. Consistent with SyRI-based comparisons at the DNA level, the annotations demonstrate a high degree of synteny while also revealing differences in structure and gene order (Figure 6). Together, these results establish differences between Pavon and IWGSC RefSeq v2.1 at the DNA and gene annotation levels and demonstrate that they may differ in biologically relevant ways, necessitating cultivar-specific references for accurate experimental analysis.

**Figure 6.**
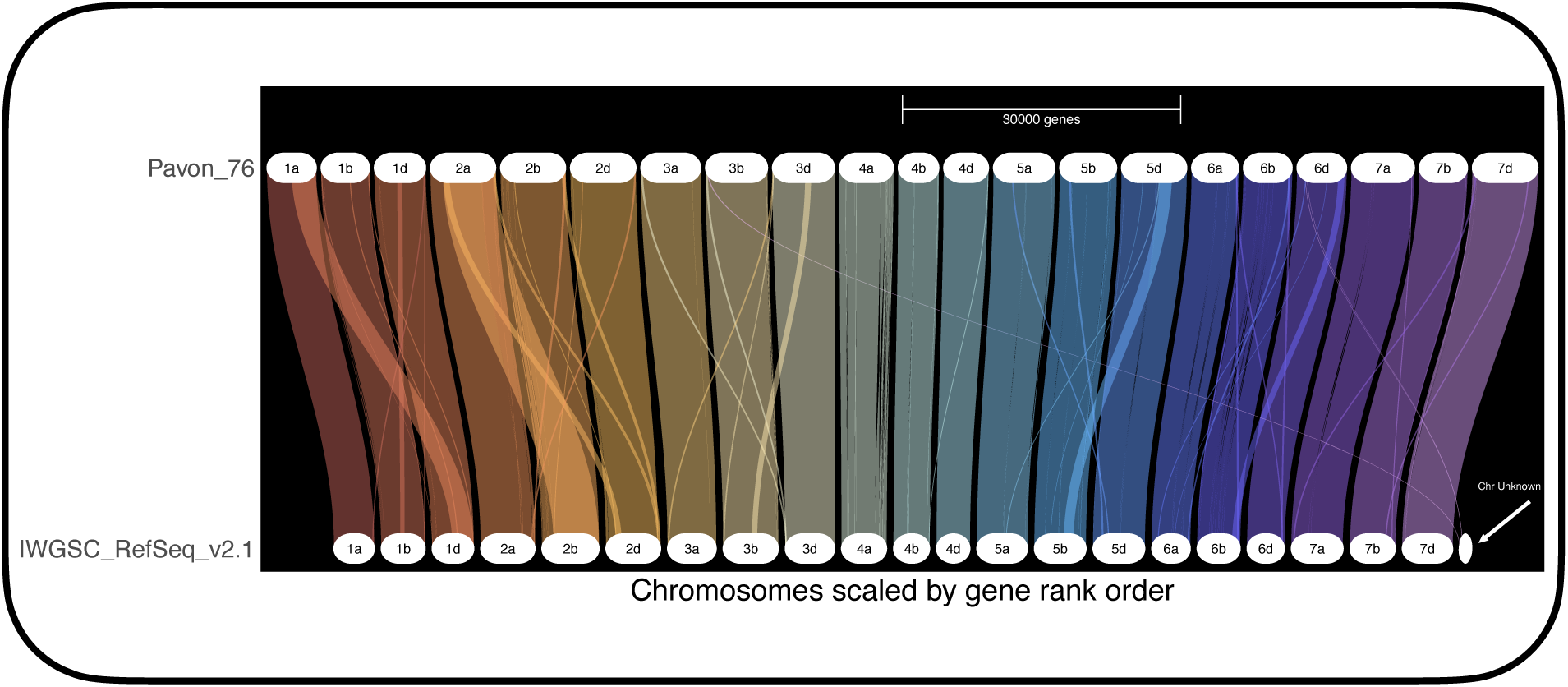
GeneSpace Results Comparing the Pavon 76 and IWGSC RefSeq v2.1 (Chinese Spring) annotations.

## 4. Discussion

This study presents a high-quality *de novo* assembly of *Triticum aestivum* cv. Pavon 76, a globally distributed Green Revolution wheat cultivar used extensively in breeding and cytogenetic research. As in prior studies, a 30-fold coverage of HiFi reads, combined with Hi-C data, has yielded 21 single-chromosome pseudomolecules with high contiguity and completeness. The evaluation of this assembly revealed an N50 of over 700 Mb and near-complete identification of BUSCO genes, comparable to or exceeding other wheat assemblies. Coverage analysis supported the genome’s accuracy, with no large-scale coverage anomalies and a low heterozygosity consistent with an inbred genome. HiFi technology drastically reduced the number of gaps relative to earlier short-read assemblies. The combination of automated scaffolding and manual curation can resolve challenging regions and produce pseudomolecules with minimal misassemblies, supported by Hi-C data (Rhie *et al*., 2021).

A key benefit of the assembly strategy is the reduction of unresolved regions compared to the IWGSC RefSeq v2.1. The large reduction in ambiguous bases and absence of coverage spikes support the integrity of the assembly and reflect the advantages of HiFi-based assemblies. Beyond comparisons of sequencing strategies, comparisons with the IWGSC RefSeq v2.1 provide genetic context for Pavon, including structural rearrangements between the two genomes (Figure 5). While wider comparisons of genome structures relative to Chinese Spring remain incompletely characterized, structural rearrangements among wheat cultivars and breeding stocks are known (Schlegel & Schlegel, 1989). Given that Pavon has been grown worldwide and is listed in the pedigrees of many newer wheats, this assembly provides a stronger genomic framework for breeding and genetic studies.

This assembly enables proper use of over 600 genetic and cytogenetic stocks developed in Pavon, spanning a wide range of research areas. This assembly provides the genomic context needed to interpret these stocks. Previous attempts to analyze these Pavon stocks were constrained by genetic differences between Pavon and the IWGSC RefSeq v2.1 assembly of Chinese Spring, often producing uninterpretable results. In contrast, this assembly resolves these discrepancies, allowing accurate characterization of these stocks and establishing a reliable foundation for future studies. In particular, its application to studies of meiotic recombination, including the role of *Ph1* in crossover site allocation and mechanisms underlying male sterility, highlights Pavon’s broad utility. By including nuclear, mitochondrial, and chloroplast genomes, this comprehensive assembly presents a powerful resource for unleashing the full potential of Pavon-derived germplasm.

More fundamentally, this Pavon assembly fills a gap in CIMMYT-derived Green Revolution wheat genome assemblies. The Green Revolution was essential to food security in developing regions by introducing wheat varieties with superior agricultural traits, such as higher yields and resistance to pests and environmental stresses, including drought (John & Babu, 2021). Despite the global influence of CIMMYT germplasm, which is grown across 60 million hectares globally (CIMMYT, n.d.), there is no comparable *de novo* assembly of a CIMMYT wheat (Liu *et al*., 2026). This Pavon assembly may serve as a representative of other CIMMYT-derived wheats and provides a foundation for future functional genomics and crop improvement studies. Our findings highlight the intrinsic limitations of relying on a single reference genome and demonstrate how biologically meaningful structural variation across wheat lineages can be obscured without cultivar-specific, *de novo* assemblies.

## 5. Conclusion

This Pavon assembly provides a high-quality wheat reference genome and annotation, including 21 nuclear pseudomolecules, mitochondrial, and chloroplast genomes, for a wheat cultivar that has been central to cytogenetic and breeding research for decades. Beyond its technical quality, this work fills a critical gap, as no *de novo* chromosome-scale assembly of a CIMMYT-derived wheat cultivar has ever been reported, despite its worldwide influence. By providing genomic context for over 600 Pavon-derived genetic stocks, this assembly will support future studies in functional genomics, wheat breeding, and crop improvement. Broadly, these results establish that reliance on a single reference genome can obscure biologically meaningful variation and underscore the need for cultivar-specific assemblies to understand wheat genome diversity.

## 6. Acknowledgments

This work was supported in part by funding from the United States Department of Agriculture, National Institute of Food and Agriculture, USDA - NIFA #CA-R-BPS-5411-H to Adam J. Lukaszewski. This material is based upon work that is supported by the National Institute of Food and Agriculture, U.S. Department of Agriculture, under award numbers 2023-70029-41306 and 2022-70029-38507. Additionally, this publication was supported by the Specialty Crop Multi-State Program from the U.S. Department of Agriculture’s (USDA) Agricultural Marketing Service in partnership with the California Department of Food and Agriculture (CDFA), under Award Number AM19SCMPCA0002-01. Joanna Melonek acknowledges support from the Australian Research Council Future Fellowship grant (FT220100792) and the Australian National University 2.0 Futures Award.

We thank Professor Adam J. Lukaszewski, University of California, Riverside, for his guidance throughout the project, including providing essential background information, the tissue of Pavon 76, and manuscript preparation. His contributions were essential to this work. We thank Professor Martin Mascher, Leibniz Institute of Plant Genetics and Crop Plant Research, and Professor Mingcheng Luo, University of California, Davis, for their advice regarding bioinformatic approaches for assembly and annotation. We also thank Pauline Lasserre-Zuber, INRAE-UCA 1095 GDEC, Clermont-Ferrand, France, and Helene Rimbert, INRAE, French National Research Institute for Agriculture, Food and Environment, GDEC, Genetics, Diversity and Ecophysiology of Cereals Joint Research Unit 5 Chemin de Beaulieu, 63000 Clermont-Ferrand, France, for their advice and support for the annotation process.

## 8. Author Contributions

EE and ZJ designed, planned, and supervised the project. MR extracted HMW DNA, and WZ prepared the PacBio HiFi libraries and facilitated PacBio HiFi sequencing. EE assembled, annotated, and analyzed the genome with guidance and intellectual contributions from ZJ, GL, JC, LW, ZS, and JM. Bioinformatic analyses were conducted by EE, GL, QL, FN, RC, TW, and SL. All authors revised, read, and approved the final manuscript.

## 9. Data Availability

The genome sequence data generated and analyzed during this study have been deposited to the National Center for Biotechnology Information (NCBI) under BioProject accession number PRJNA1464003. Raw PacBio HiFi sequencing reads and Hi-C sequencing reads are available through the Sequence Read Archive (SRA) under accessions SRR38739884 and SRR38739883, respectively. Individual sample metadata are available under BioSample accession number SAMN59652007. The assembled genomes have been submitted to GenBank under accession number JBYCRE000000000. All sequence datasets are publicly accessible and can be retrieved via the NCBI website (http://www.ncbi.nlm.nih.gov/bioproject/1464003).

