## Supplemental Figure 1 for "Genome Assembly of the Green Revolution Wheat Cultivar Pavon 76 Establishes a Reference for CIMMYT-Derived Wheat"

### Supplemental Figure 1. NucFreq Analysis of Pavon 76 by Chromosome (Chr 1A - 3A)

#### Chr 1A

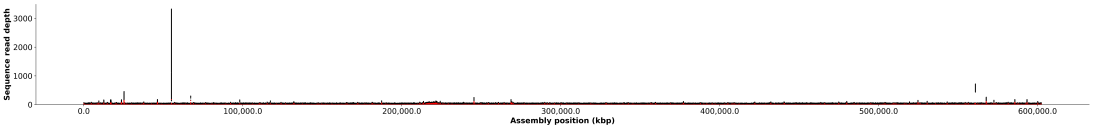


#### Chr 1B

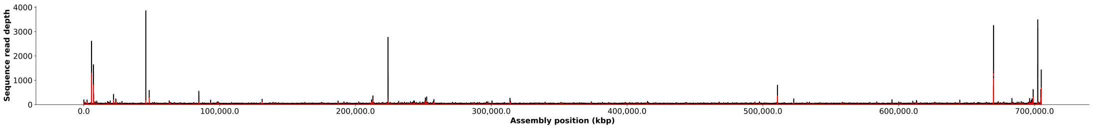


#### Chr 1D

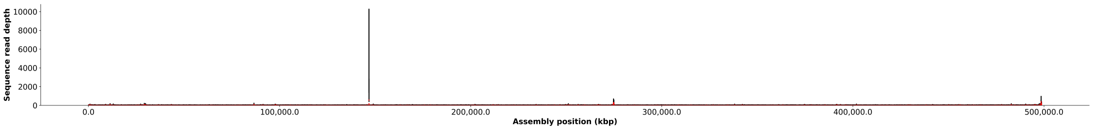


#### Chr 2A

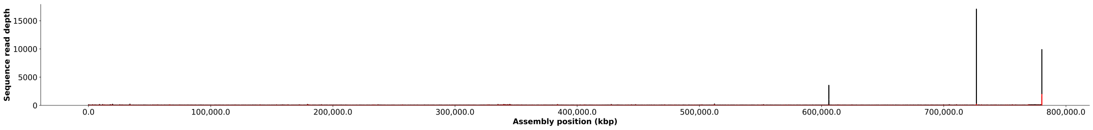


#### Chr 2B

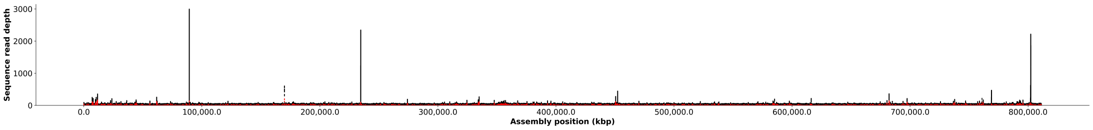


#### Chr 2D

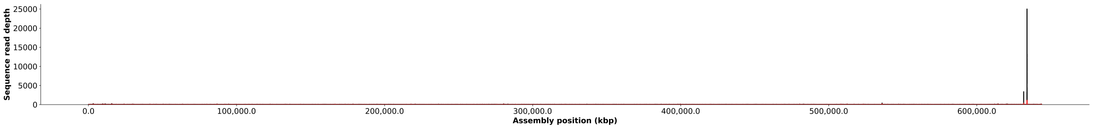


#### Chr 3A

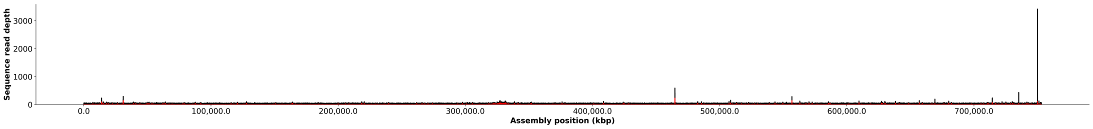

### Supplemental Figure 1. NucFreq Analysis of Pavon 76 by Chromosome (3B - 5B)

#### Chr 3B

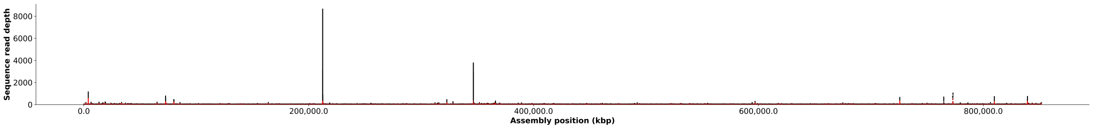


#### Chr 3D

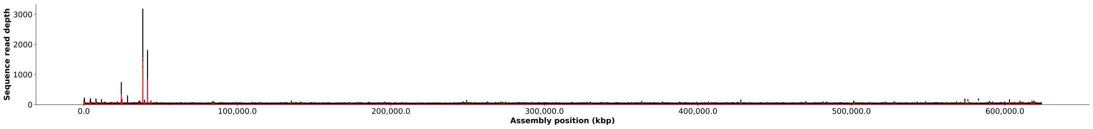


#### Chr 4A

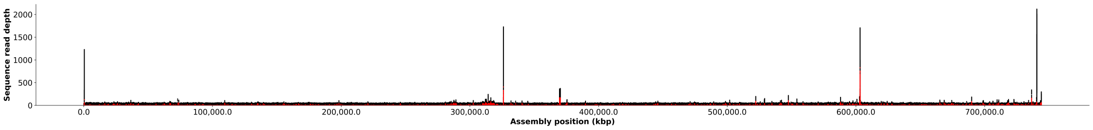


#### Chr 4B

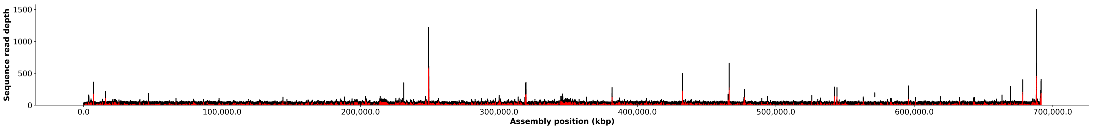


#### Chr 4D

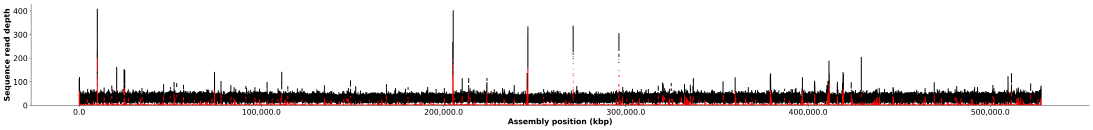


#### Chr 5A

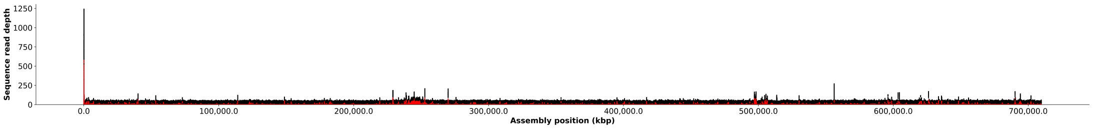


#### Chr 5B

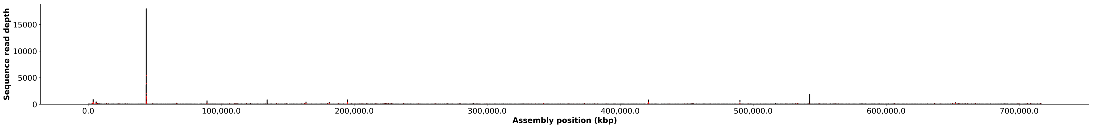

### Supplemental Figure 1. NucFreq Analysis of Pavon 76 by Chromosome (5D - 7D)

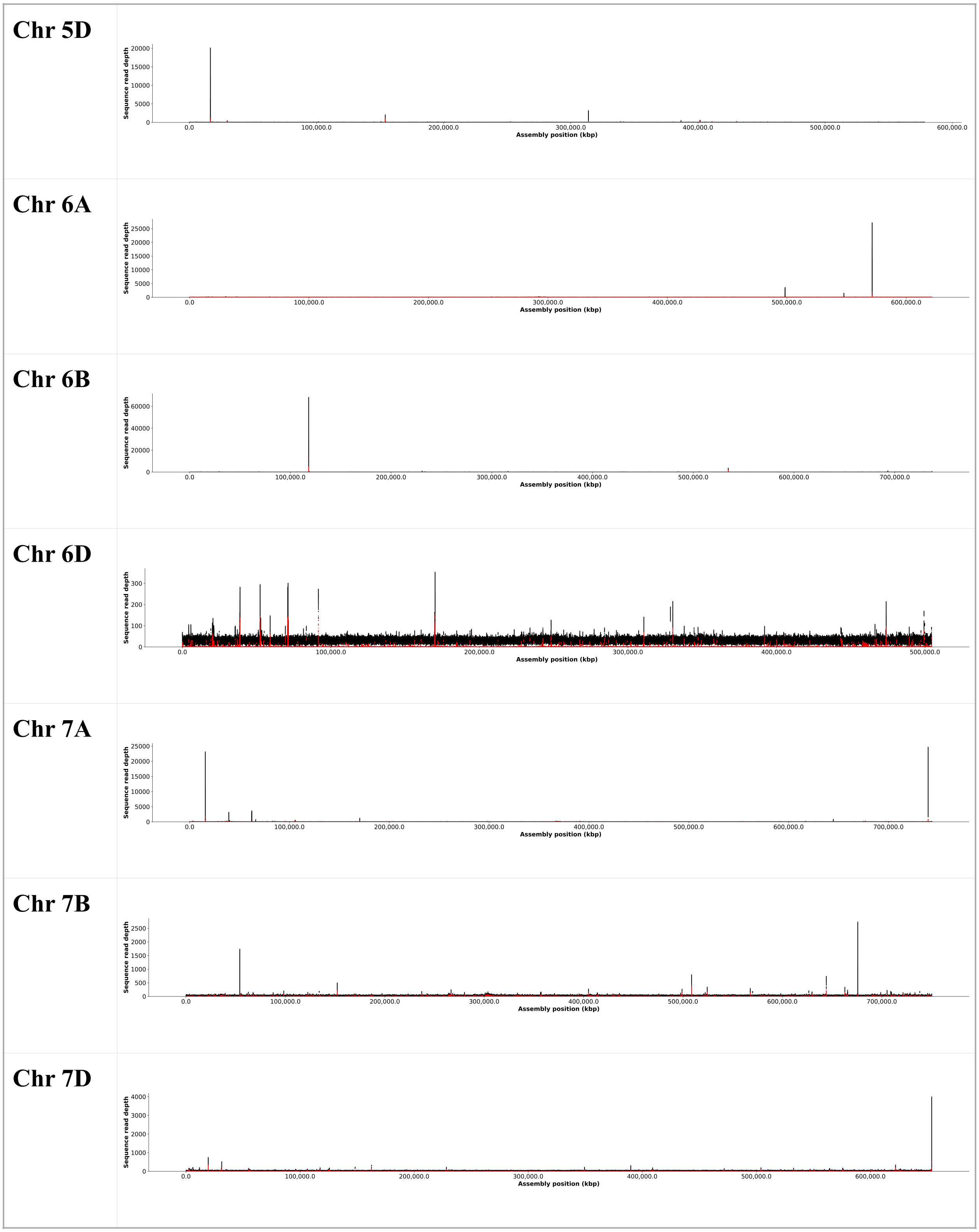
