## Supplemental Figure 2 for "Genome Assembly of the Green Revolution Wheat Cultivar Pavon 76 Establishes a Reference for CIMMYT-Derived Wheat"

**Supplemental Figure 2. NucFreq Analysis of Pavon 76 by Chromosome (Chr 1A - 3A) with Sequence Read Depth Limit of 100**

#### Chr 1A

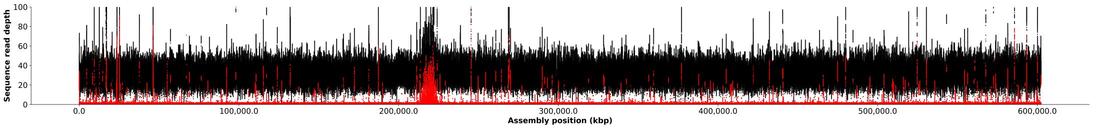


#### Chr 1B

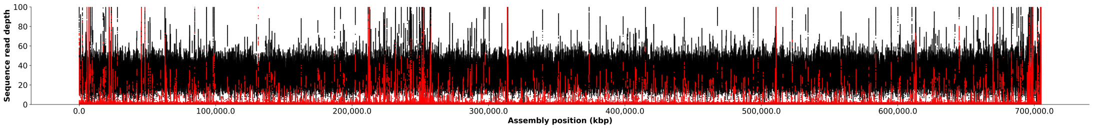


#### Chr 1D

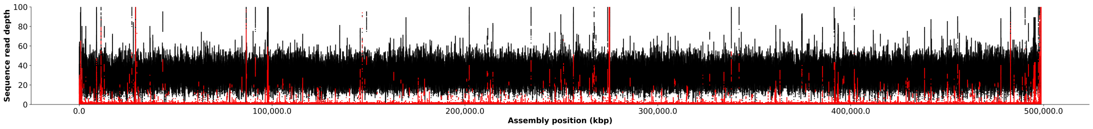


#### Chr 2A

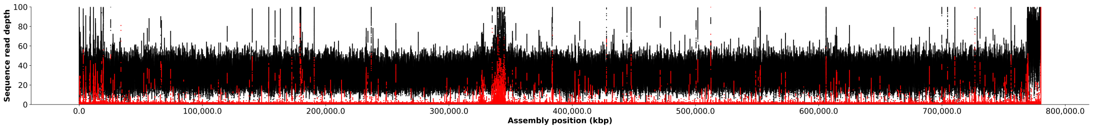


#### Chr 2B

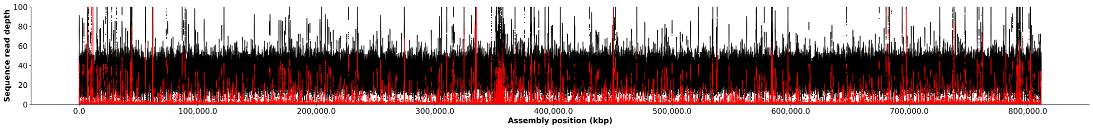


#### Chr 2D

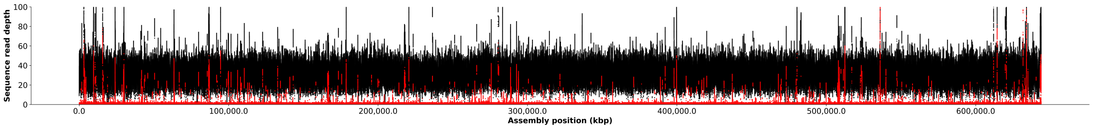


#### Chr 3A

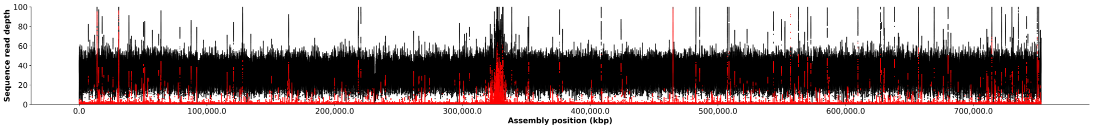

### Supplemental Figure 2. NucFreq Analysis of Pavon 76 by Chromosome (Chr 3B - 5B) with Sequence Read Depth Limit of 100

#### Chr 3B

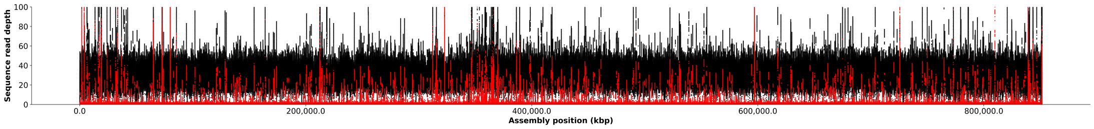


#### Chr 3D

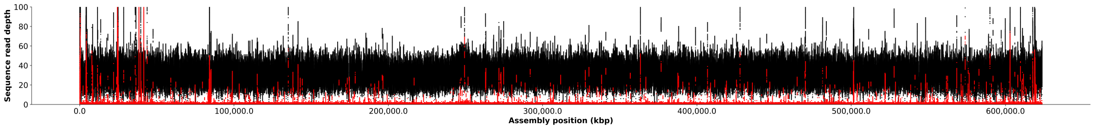


#### Chr 4A

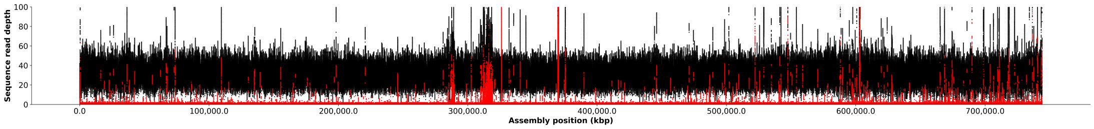


#### Chr 4B

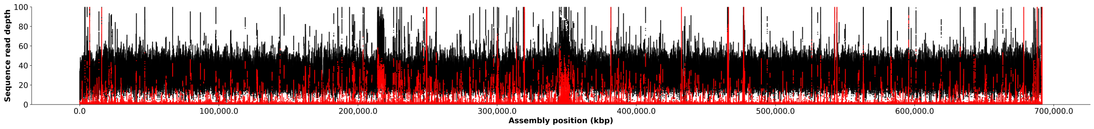


#### Chr 4D

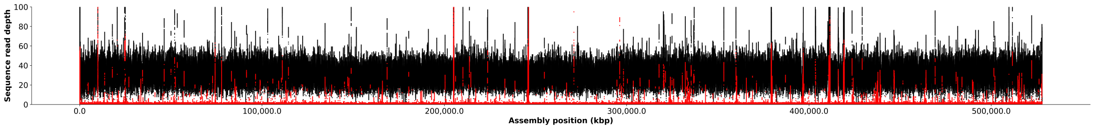


#### Chr 5A

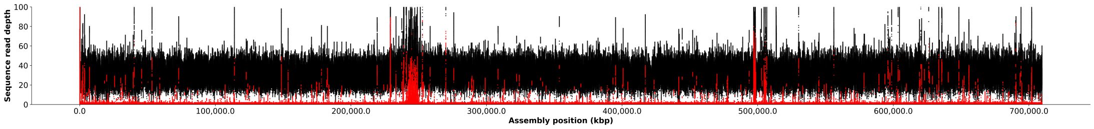


#### Chr 5B

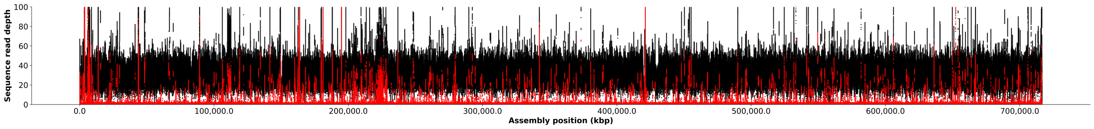

### Supplemental Figure 2. NucFreq Analysis of Pavon 76 by Chromosome (Chr 5D - 7D) with Sequence Read Depth Limit of 100

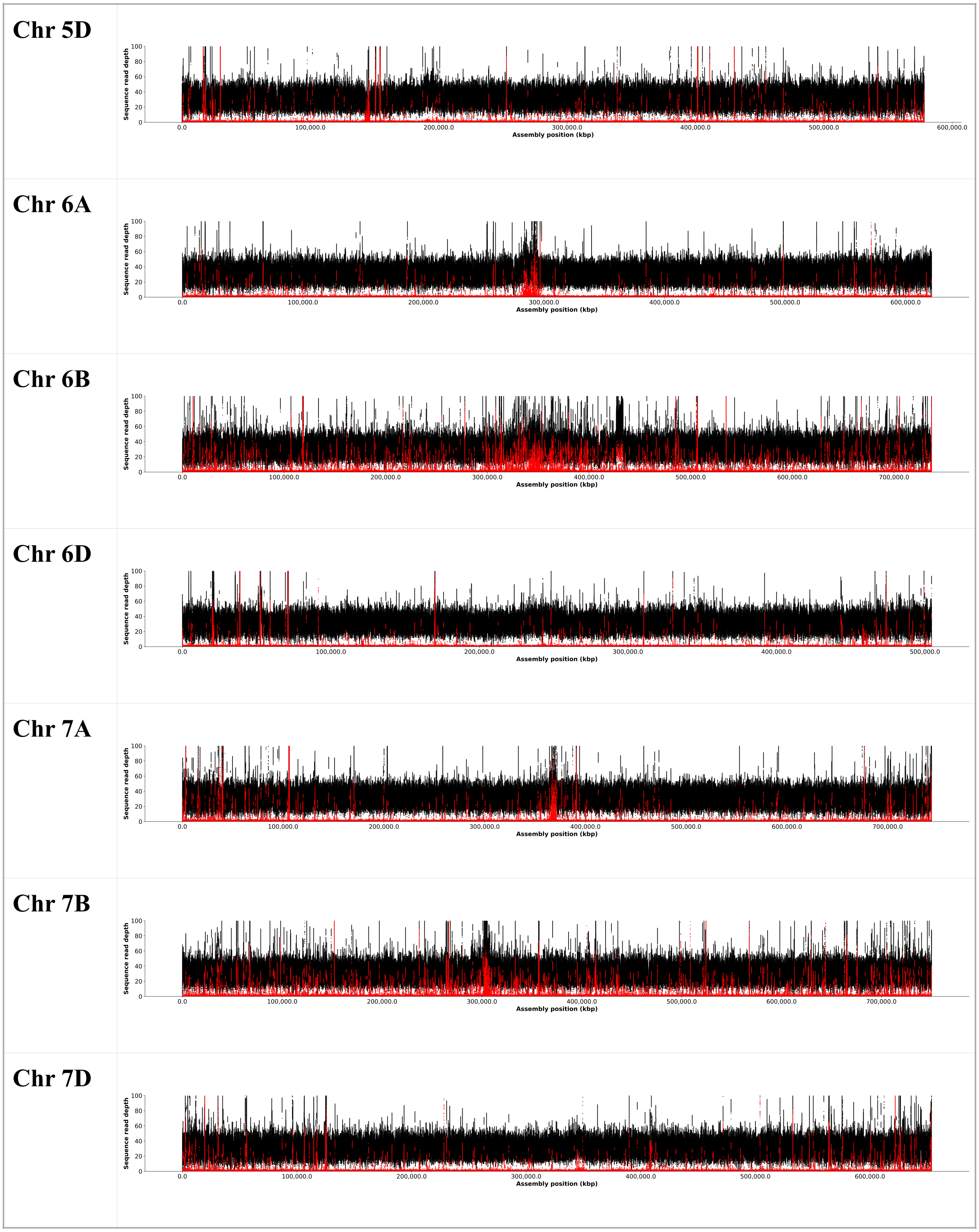
