## Supplemental Table 1 for "Genome Assembly of the Green Revolution Wheat Cultivar Pavon 76 Establishes a Reference for CIMMYT-Derived Wheat"

**Supplemental Table 1. Pavon 76 CCS Analysis Report Summary Metrics per SMRT Cell**

| | **m84066_250516_203254_s2** | **m84066_250522_212353_s2** | **m84066_250529_191935_s1** | **m84066_250529_212203_s2** | **m84066_250529_232437_s3** | **m84066_250609_193946_s1** |
|---|---|---|---|---|---|---|
| HiFi reads | 7.6 M | 7.6 M | 5.9 M | 6.3 M | 6.2 M | 6.5 M |
| HiFi reads yield | 90.61 Gb | 92.01 Gb | 70.36 Gb | 74.75 Gb | 74.70 Gb | 78.68 Gb |
| HiFi reads length (mean) | 11.95 kb | 12.04 kb | 12.02 kb | 11.94 kb | 11.95 kb | 12.17 kb |
| HiFi reads length (median, bp) | 11,344 | 11,433 | 11,422 | 11,351 | 11,356 | 11,568 |
| HiFi Read Length N50 (bp) | 12,032 | 12,135 | 12,092 | 12,006 | 12,021 | 12,268 |
| HiFi Read Quality (median) | Q39 | Q40 | Q39 | Q39 | Q40 | Q40 |
| HiFi Read Quality (median) | 39 | 40 | 39 | 39 | 40 | 40 |
| Base Quality ≥Q30 (%) | 95.90% | 96.04% | 95.92% | 95.82% | 95.94% | 96.08% |
| HiFi Number of Passes (mean) | 12 | 12 | 11 | 12 | 12 | 12 |
| Missing adapters (%) | 2.36% | 2.38% | 2.38% | 2.34% | 2.33% | 2.45% |
