## Supplemental Table 2 for "Genome Assembly of the Green Revolution Wheat Cultivar Pavon 76 Establishes a Reference for CIMMYT-Derived Wheat"

**Supplemental Table 2. Gene Composition of the Mitogenome of Pavon 76**

| Group of Genes Encoding Subunits of: | Gene Name |
|---|---|
| Complex I (NADH dehydrogenase) | *nad1(×1)***, *nad2(×1)***, *nad3*, *nad4****, *nad4L*, *nad5(×1)**, *nad6*, *nad7 *****, *nad9* |
| Complex III (ubiquinol-cytochrome c reductase) | *cob(×1)* |
| Complex IV (cytochrome c oxidase) | *cox1*, *cox2 (×1)**, *cox3* |
| Complex V (ATP synthase) | *atp1(×4)*, *atp4(×2)*, *atp6***(×2)***, *atp8***(×2,** ×2**)***, *atp9(×2)* |
| Ribosomal proteins (SSU) | *rps1(×1)*, *rps2*, *rps3*, *rps4*, *rps7*, *rps12(×1)*, *rps13*, *rps19(×1)* |
| Ribosomal proteins (LSU) | *rpl5*, *rpl16* |
| Maturases | *matR* |
| Transfer RNAs (tRNAs) | *trnA **, *trnC*, *trnD***(×2)***, *trnE*, *trnF*, *trnfM***(×3)***, *trnH*, *trnK***(×3)***, *trnL*, *trnM***(×4)***, *trnN*, *trnP(×***2***)*, *trnQ(×***3***)*, *trnS(×***3***)*, *trnW*, *trnY* |
| Ribosomal RNAs (rRNAs) | *rrn5(×***3***)*, *rrn18(×***3***)*, *rrn26(×***3***)* |
| Other genes | *ccmB*, *ccmC*, *ccmFC*, *ccmFN*, *mttB* |

Note: * The asterisks denote the number of introns (**, ***, **** indicate two, three, four introns, respectively); the numbers in parentheses represent the number of additional copies; bold numbers indicate additional full-length copies, whereas non-bold numbers indicate partial-length copies.
