## Supplemental Table 3 for "Genome Assembly of the Green Revolution Wheat Cultivar Pavon 76 Establishes a Reference for CIMMYT-Derived Wheat"

**Supplemental Table 3. Gene Functions of the Chloroplast Genome of Pavon 76**

| Category for function | Function groups for genes | Gene |
|---|---|---|
| Photosynthesis | Photosystem I | *psaA(×1), psaB(×1), psaC, psaI, psaJ* |
| | Photosystem II | *psbA, psbB, psbC, psbD, psbE, psbF, psbH, psbI, psbJ, psbK, psbL, psbM, psbT, psbZ* |
| | ATP synthase | *atpA, atpB, atpE, atpF*, atpH, atpI* |
| | ATP-dependent protease subunit P gene | *clpP1** |
| | Cytochrome b/f complex | *petA, petB*, petD*, petG, petL, petN* |
| | Large subunit of rubisco | *rbcL* |
| | NADH dehydrogenase | *ndhA*, ndhB (**×2**)**, ndhC, ndhD, ndhE, ndhF, ndhG, ndhH(×1), ndhI, ndhJ, ndhK* |
| Expression | RNA polymerase | *rpoA, rpoB, rpoC1*, rpoC2* |
| | Proteins of small ribosomal subunit (SSU) | *rps2, rps3(**×4**), rps4, rps7(**×2**), rps8, rps11, rps12**, rps14, rps15(**×2**), rps16*, rps18, rps19(**×2**)* |
| | Proteins of large ribosomal subunit (LSU) | *rpl2**, rpl14, rpl16, rpl20, rpl22, rpl23(**×2,** ×1), rpl32, rpl33, rpl36* |
| | Transfer RNAs | *trnA-UGC(**×4**), trnC-GCA, trnD-GUC, trnE-UUC, trnF-GAA, trnfM-CAU(**×2**), trnG-UCC, trnH-GTG(**×2**), trnI-CAU(**×2**), trnI-GAU(**×4**), trnK-UUU(**×2**), trnL-CAA(**×2**), trnL-UAA(**×2**), trnL-UAG, trnM-CAU(**×6**), trnN-GUU(**×2**), trnP-UGG, trnQ-UUG, trnR-ACG(**×2**), trnR-UCU, trnS-CGA*, trnS-GCU, trnS-GGA, trnS-UGA, trnT-GGU(**×2**), trnT-UGU, trnV-GAC(**×2**), trnV-UAC(**×2**), trnW-CCA, trnY-GUA* |
| | Ribosomal RNAs | *rrn4.5 (**×2**), rrn5 (**×2**), rrn16 (**×2**), rrn23 (**×2**)* |
| | C-type cytochrome synthesis gene | *ccsA* |

| | | |
|---|---|---|
| Others | Envelope membrane protein | *cemA* |
| | Maturase | *matK* |
| | Translation initiation factor | *infA* |
| unknown function | Conserved open reading frame | *ycf1(×**2**)**, ycf2(×**2**), ycf68(×**2**)* |

Note: * The asterisk denotes the number of introns (**, ***, **** is for two, three, four introns, respectively); the numbers in parentheses represent the number of additional copies; bold numbers indicate additional full-length copies, whereas non-bold numbers indicate partial-length copies.
