## Supplemental File 1 for "Genome Assembly of the Green Revolution Wheat Cultivar Pavon 76 Establishes a Reference for CIMMYT-Derived Wheat"

### Report

| | Triticum_Aestivum_Pavon76_Nuclear_Genome_Chromosomes |
|---|---|
| # contigs (>= 0 bp) | 21 |
| # contigs (>= 1000 bp) | 21 |
| # contigs (>= 5000 bp) | 21 |
| # contigs (>= 10000 bp) | 21 |
| # contigs (>= 25000 bp) | 21 |
| # contigs (>= 50000 bp) | 21 |
| Total length (>= 0 bp) | 14248756236 |
| Total length (>= 1000 bp) | 14248756236 |
| Total length (>= 5000 bp) | 14248756236 |
| Total length (>= 10000 bp) | 14248756236 |
| Total length (>= 25000 bp) | 14248756236 |
| Total length (>= 50000 bp) | 14248756236 |
| # contigs | 21 |
| Largest contig | 851610634 |
| Total length | 14248756236 |
| GC (%) | 46.14 |
| N50 | 709756138 |
| N90 | 527888988 |
| auN | 692065507.5 |
| L50 | 10 |
| L90 | 19 |
| # N's per 100 kbp | 0.72 |

All statistics are based on contigs of size >= 3000 bp, unless otherwise noted (e.g., "# contigs (>= 0 bp)" and "Total length (>= 0 bp)" include all contigs).

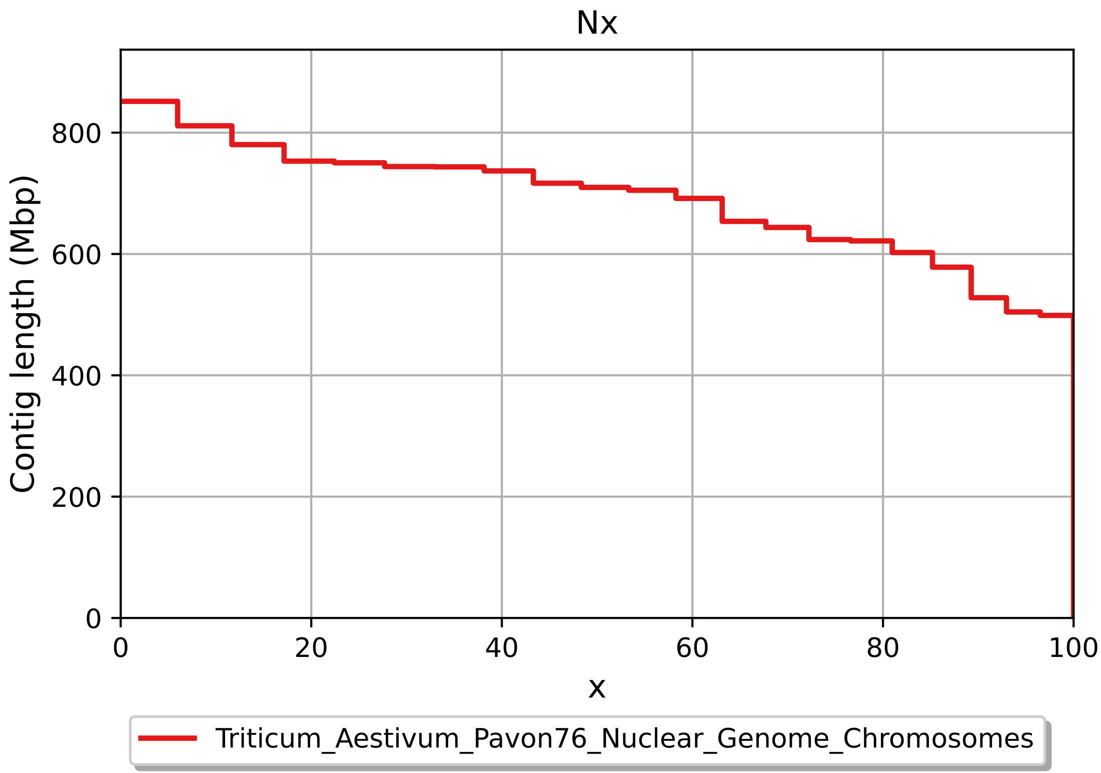

Nx
Contig length (Mbp)
0
200
400
600
800
0
20
40
60
80
100
x
Triticum_Aestivum_Pavon76_Nuclear_Genome_Chromosomes

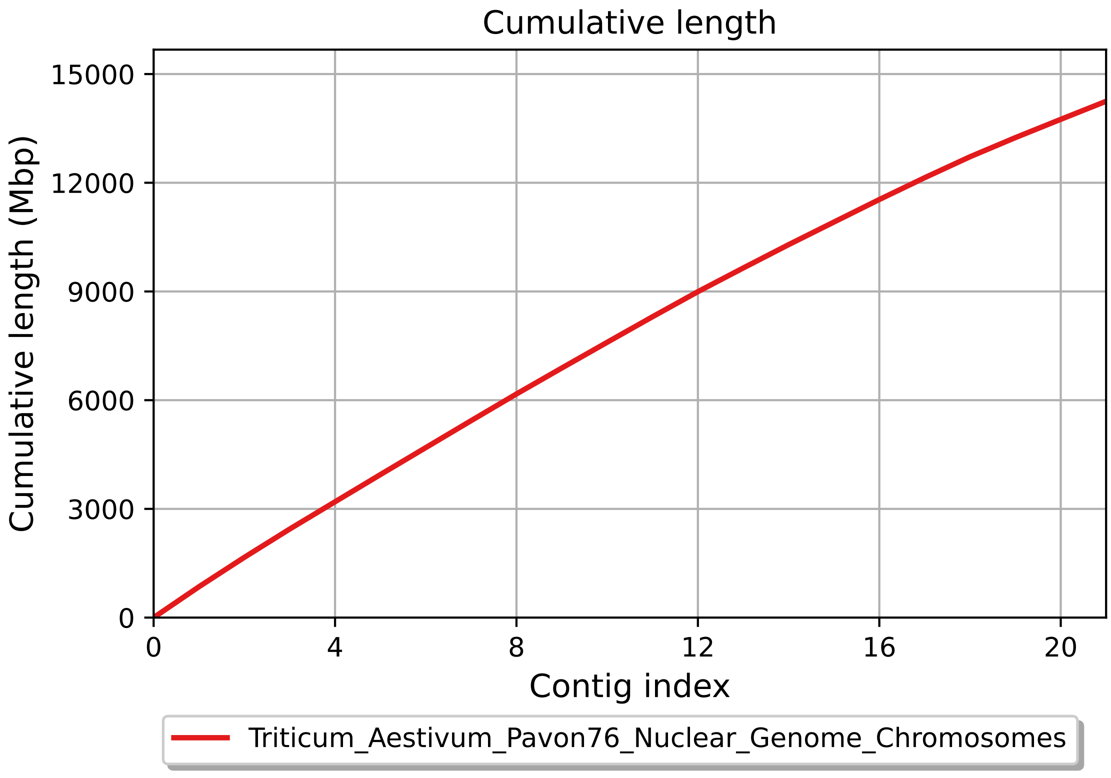
Cumulative length
Cumulative length (Mbp)
0
3000
6000
9000
12000
15000
0
4
8
12
16
20
Contig index
Triticum_Aestivum_Pavon76_Nuclear_Genome_Chromosomes

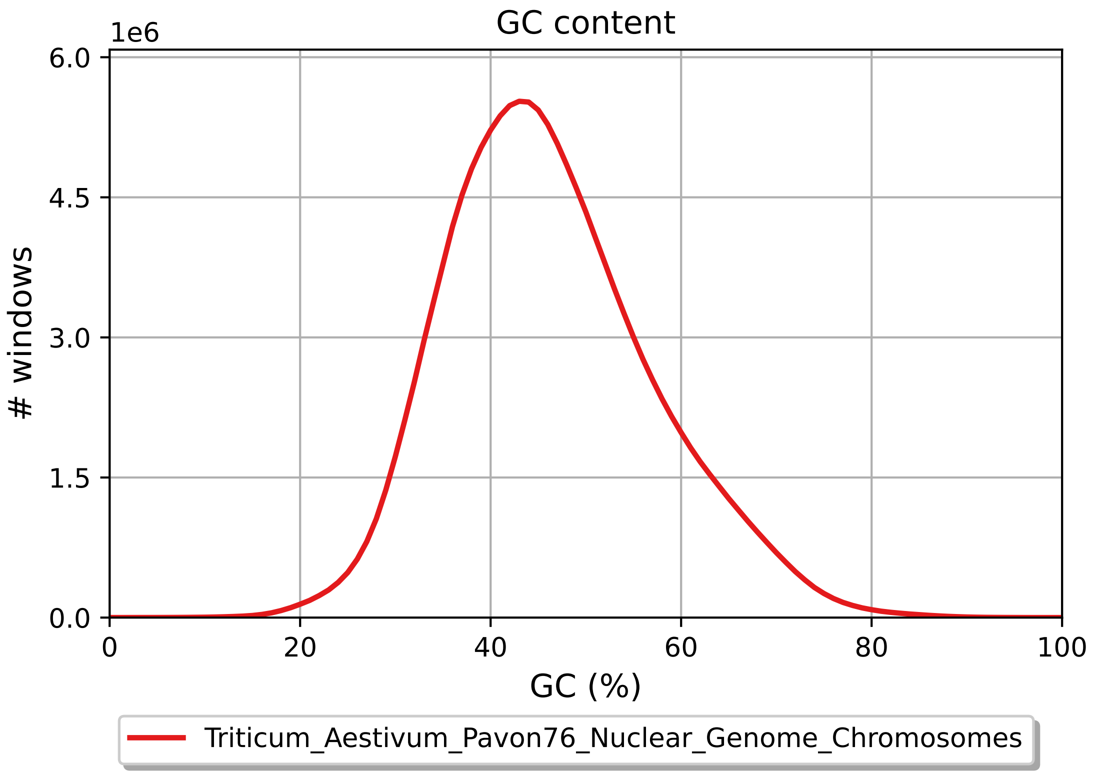

GC content
1e6
6.0
4.5
3.0
1.5
0.0
### windows
0
20
40
60
80
100
GC (%)
Triticum_Aestivum_Pavon76_Nuclear_Genome_Chromosomes

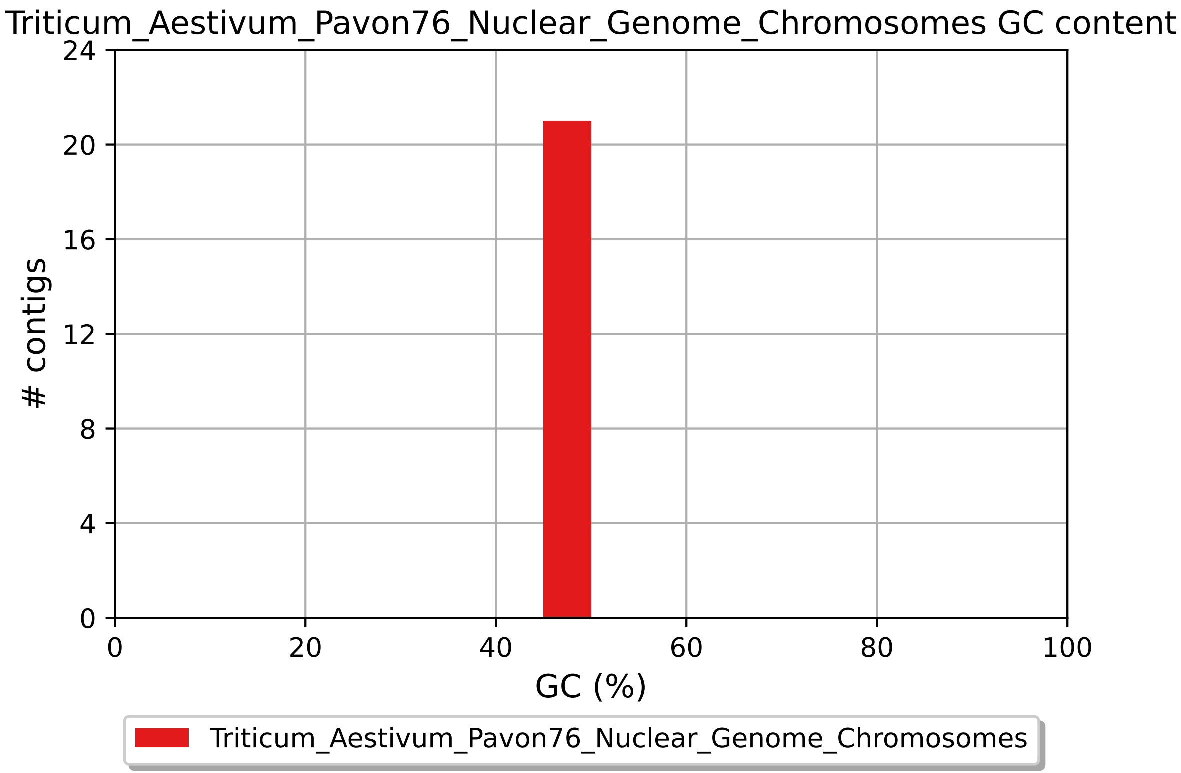

Triticum_Aestivum_Pavon76_Nuclear_Genome_Chromosomes GC content
### contigs
0
4
8
12
16
20
24
0
20
40
60
80
100
GC (%)
Triticum_Aestivum_Pavon76_Nuclear_Genome_Chromosomes
