## Supplemental File 2 for "Genome Assembly of the Green Revolution Wheat Cultivar Pavon 76 Establishes a Reference for CIMMYT-Derived Wheat"

### Report

| | iwgsc_refseqv2.1_assembly_Chromosomes_Only |
|---|---|
| # contigs (>= 0 bp) | 21 |
| # contigs (>= 1000 bp) | 21 |
| # contigs (>= 5000 bp) | 21 |
| # contigs (>= 10000 bp) | 21 |
| # contigs (>= 25000 bp) | 21 |
| # contigs (>= 50000 bp) | 21 |
| Total length (>= 0 bp) | 14225829371 |
| Total length (>= 1000 bp) | 14225829371 |
| Total length (>= 5000 bp) | 14225829371 |
| Total length (>= 10000 bp) | 14225829371 |
| Total length (>= 25000 bp) | 14225829371 |
| Total length (>= 50000 bp) | 14225829371 |
| # contigs | 21 |
| Largest contig | 851934019 |
| Total length | 14225829371 |
| GC (%) | 46.17 |
| N50 | 713360525 |
| N90 | 518332611 |
| auN | 691971349.3 |
| L50 | 10 |
| L90 | 19 |
| # N's per 100 kbp | 1524.94 |

All statistics are based on contigs of size >= 3000 bp, unless otherwise noted (e.g., "# contigs (>= 0 bp)" and "Total length (>= 0 bp)" include all contigs).

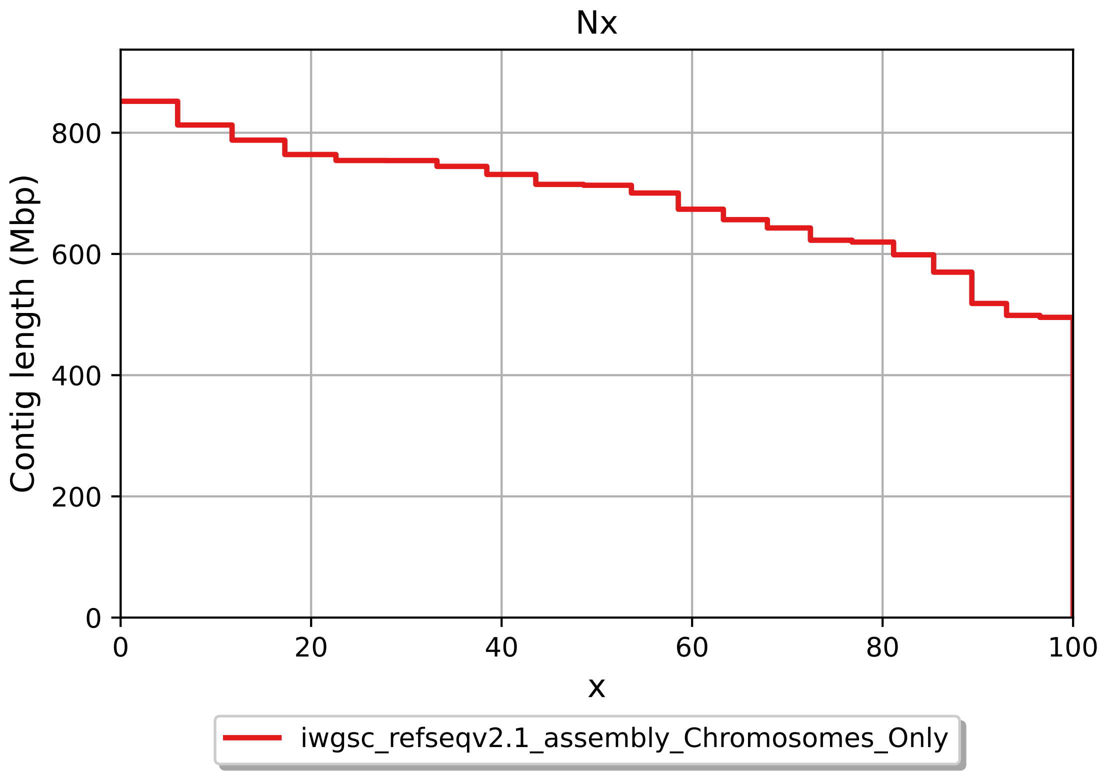
Nx
Contig length (Mbp)
0
200
400
600
800
0
20
40
60
80
100
x
iwgsc_refseqv2.1_assembly_Chromosomes_Only

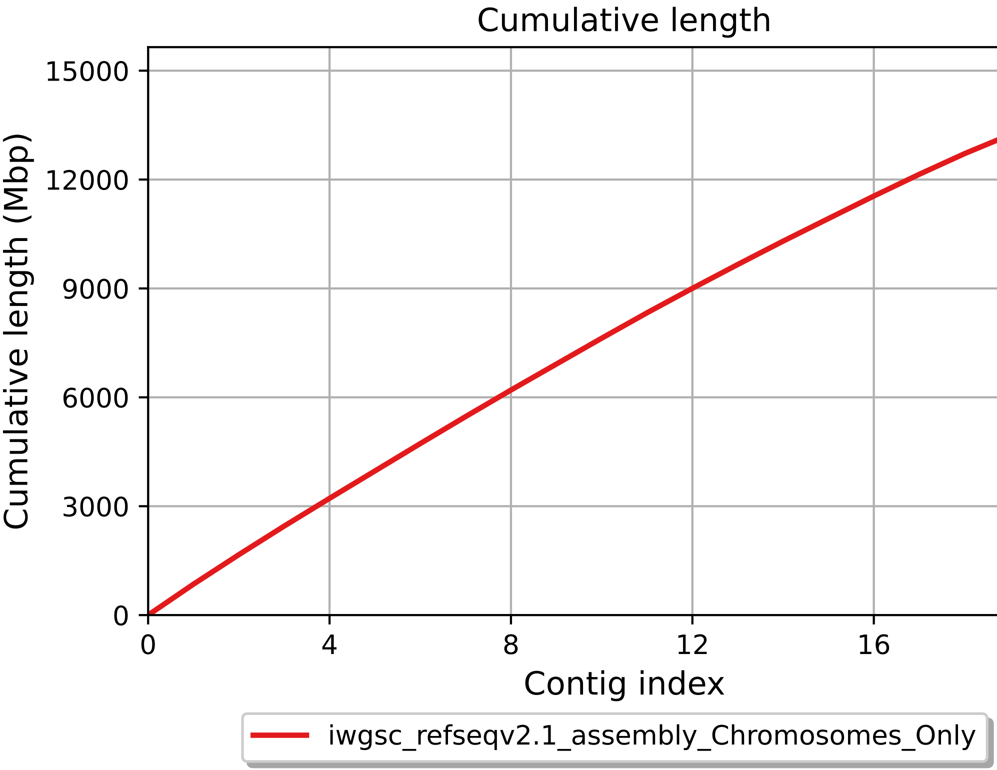
Cumulative length
Cumulative length (Mbp)
0
3000
6000
9000
12000
15000
0
4
8
12
16
20
Contig index
iwgsc_refseqv2.1_assembly_Chromosomes_Only

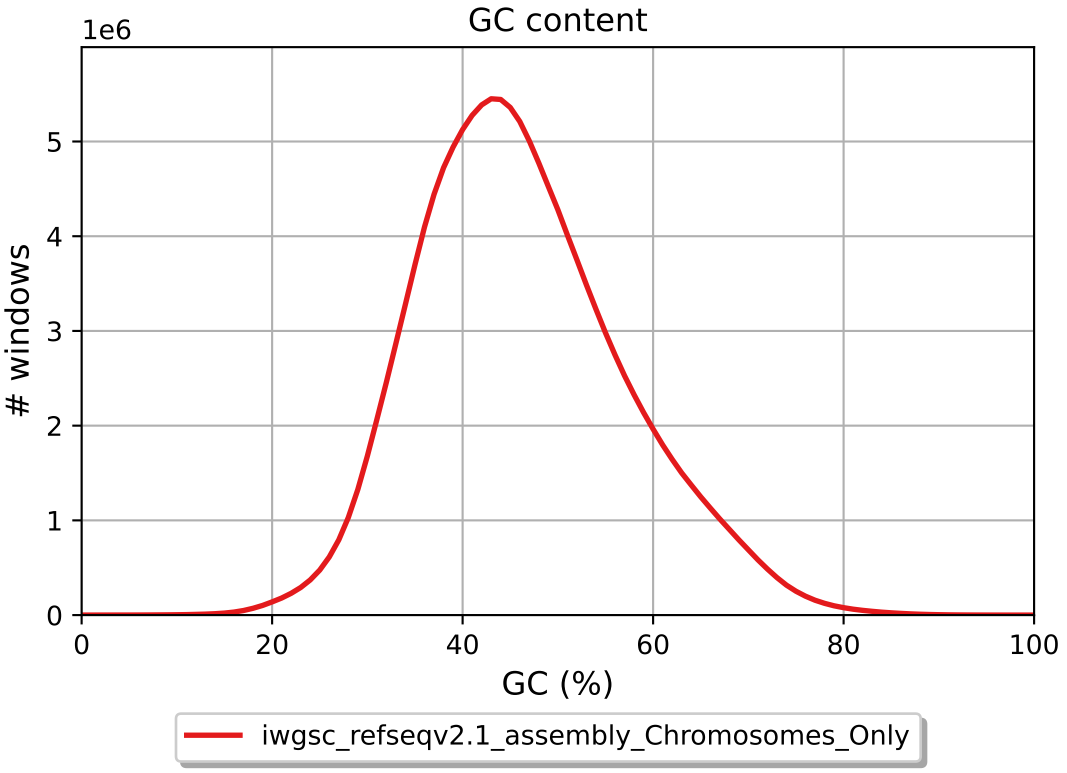
GC content
1e6
### windows
0
1
2
3
4
5
0
20
40
60
80
100
GC (%)
iwgsc_refseqv2.1_assembly_Chromosomes_Only

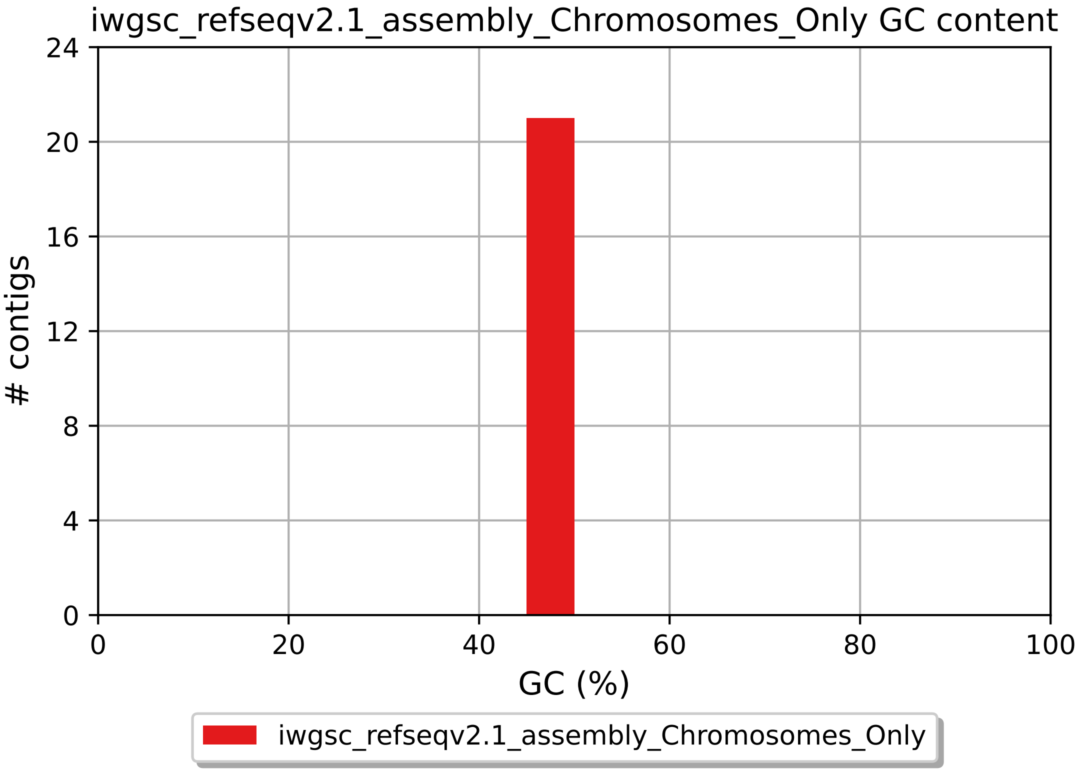
iwgsc_refseqv2.1_assembly_Chromosomes_Only GC content
### contigs
0
4
8
12
16
20
24
0
20
40
60
80
100
GC (%)
iwgsc_refseqv2.1_assembly_Chromosomes_Only
